# Interrogating the structure and function of the human voltage-gated proton channel (hHv1) with a fluorescent noncanonical amino acid

**DOI:** 10.1101/2025.10.23.684235

**Authors:** Emerson M. Carmona, William N. Zagotta, Sharona E. Gordon

## Abstract

The human voltage-gated proton channel (hH_v_1) is a dimer of voltage-sensor domains (VSDs) containing highly selective proton permeation pathways in each monomer. In addition to voltage, hH_v_1 is regulated by other stimuli, including pH gradients, mechanical forces, and ligands such as Zn^2+^. Aside from the VSDs, this membrane protein contains an N-terminal domain and a C-terminal coiled-coil domain (CC) formed between the monomers. To address the need for direct measurements of conformational rearrangements in hH_v_1, we developed a Förster resonance energy transfer (FRET) approach to measuring the conformational rearrangements in full-length hH_v_1 purified from *E. coli*. We used genetic code expansion (GCE) to generate a library of 14 full-length hH_v_1 constructs, each incorporating the fluorescent noncanonical amino acid acridon-2-ylalanine (Acd) at a different site throughout the various structural domains. Following the expression and purification of these hH_v_1-Acd proteins, we found that 12 sites yielded stable and functional proton-permeable channels. The fluorescence properties of Acd at each site showed small site-specific differences. Furthermore, we measured site-specific FRET efficiencies from tryptophan (Trp) and tyrosine (Tyr) to Acd in the hH_v_1-Acd proteins and found results consistent with correct folding in detergent micelles. Finally, the addition of Zn^2+^ produced reversible changes in FRET, with affected residues clustered on the intracellular side of the channel.

## Introduction

The human voltage-gated proton channel (hH_v_1) is a membrane protein that forms a proton selective channel (1, 2). Despite its physiological importance, its molecular mechanisms are still poorly understood, including the precise structural identity of the permeation pathway and the gating mechanism of voltage (3, 4), pH gradients (3, 5), membrane stretch (6), and ligands such as the classical inhibitor Zn^2+^ (7, 8). Understanding these processes requires connecting the functional properties to the hH_v_1 structure.

Each monomer of hH_v_1 dimers includes an intracellular N-terminal domain of unknown function, a transmembrane voltage-sensor domain (VSD), and an intracellular coiled-coil (CC), which forms the primary intersubunit interface (9, 10). Current structural models, however, have been obtained from truncated or chimeric channel constructs (11, 12), and thus, it is not clear whether they correspond to physiological or functional conformations. Moreover, the multiple kinetic components of H_v_1 gating currents suggest that the protein’s resting state comprises an ensemble of conformations (4, 5), which is consistent with the conformational flexibility inferred from structural approaches (12–14). Because protein function arises from this distribution of conformational states (15), approaches to directly measure structural heterogeneity in the full-length hH_v_1 are needed.

An attractive approach to interrogate the structural heterogeneity of hH_v_1 is fluorescence spectroscopy, which offers high sensitivity at nanomolar protein concentrations and compatibility with near-physiological conditions. Fluorescence can report on local environment and accessibility changes, as well as quantify distance changes through Förster resonance energy transfer (FRET) (16). Importantly, time-resolved FRET measured using lifetimes preserves information about conformational heterogeneity on the nanosecond timescale, which is lost in steady-state measurements due to the averaging process (17). Together, these capabilities make fluorescence, especially time-resolved FRET, a powerful method to connect hH_v_1 structure and function (18).

A few obstacles remain to directly measuring conformational heterogeneity of hH_v_1. First, a method to express and purify stable and functional protein is needed, which is challenging for membrane proteins. Second, site-specific fluorophore labeling is difficult at buried sites, resulting in low labeling efficiency. Third, traditional fluorophores are bulky and have long linkers (14), resulting in protein structure perturbation and heterogeneity dominated by the properties of the label (19). A recent method was optimized for robust expression and purification of full-length, functional hH_v_1 in *E. coli* (20, 21). The remaining obstacles can be overcome by labeling the protein with the fluorescent noncanonical amino acid acridon-2-ylalanine (Acd) using genetic code expansion (GCE) (22–24). This technology allows site-specific labeling during translation by reassigning an amber stop codon using an orthogonal aminoacyl-tRNA synthetase (RS)/tRNA (25, 26) (Figure 1A). Acd’s small size and minimal linker reduce perturbations and heterogeneity, and its favorable photophysics support both steady-state and time-resolved measurements (27). Here, we incorporated Acd across all hH_v_1 structural domains and purified 12 of 14 constructs as stable, functional proteins. We then used Acd’s environmental sensitivity and FRET from tryptophan (Trp)/tyrosine (Tyr) to Acd to validate the protein folding and report Zn^2+^-dependent conformational changes. These data establish a platform for interrogating conformational heterogeneity and dynamics in the full-length hH_v_1.

**Figure 1.**
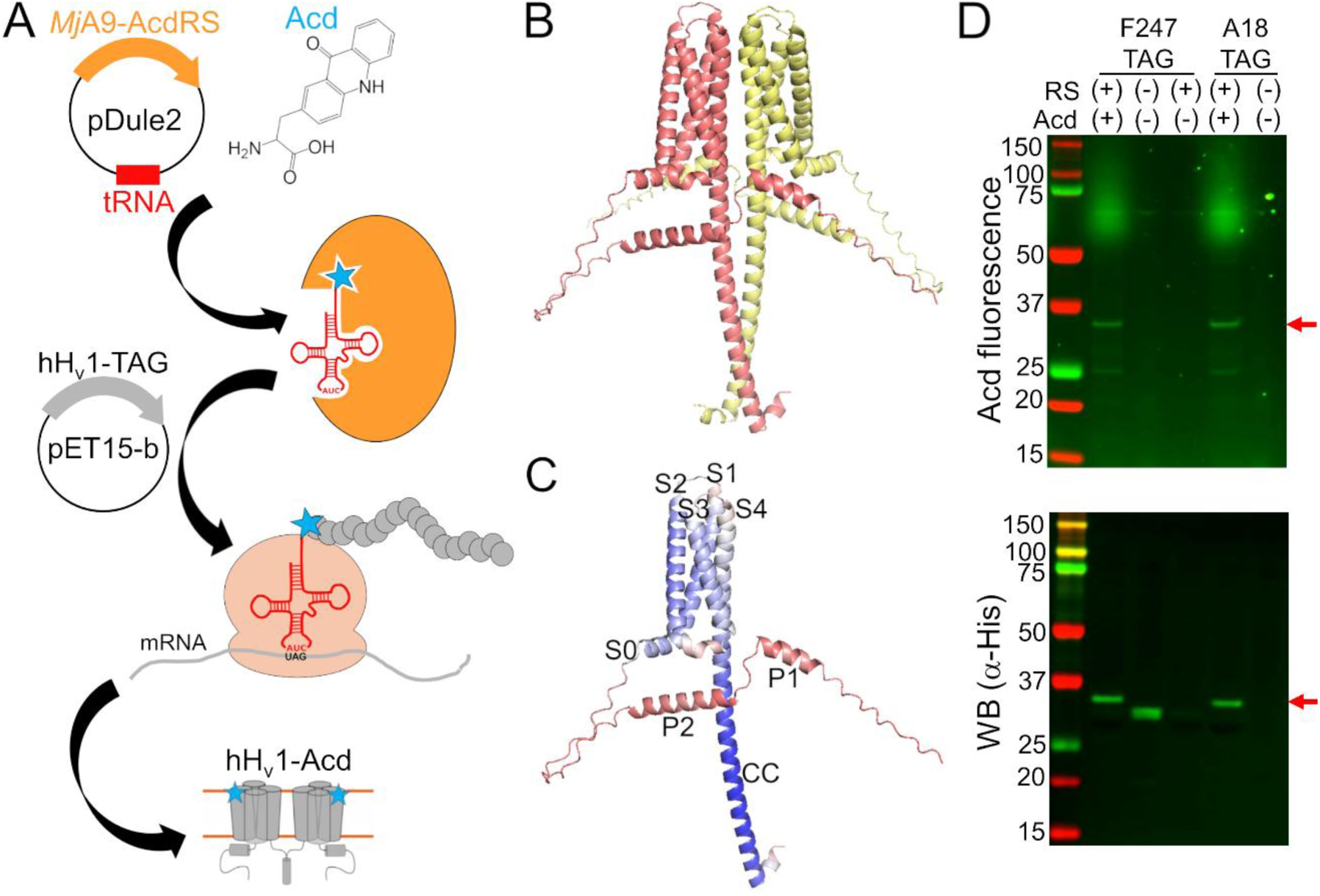
hH_v_1 was labeled with the fluorescent noncanonical amino acid Acd by genetic code expansion. (A) Cartoon showing the components of the genetic code expansion system used to incorporate Acd via amber codon suppression in hH_v_1. (B) AlphaFold structural model of the dimer hH_v_1 colored by subunit. (C) AlphaFold structural model of a single hH_v_1 subunit colored by the predicted Local Distance Difference Test (pLDDT). Model confidence was represented by a gradient from red to blue, indicating low to high pLDDT. (D) In-gel fluorescence and Western blot to evaluate Acd incorporation. An Acd fluorescence band corresponding to the molecular weight of full-length hH_v_1 (red arrow) was observed in cellular extracts after separation by SDS-PAGE only when cells contained the *Mj*A9-AcdRS/tRNA plasmid (RS) and were grown in the presence of 1mM Acd (top). The Western blot against the His-tag present at the N-terminus of hH_v_1 confirmed that the band corresponded to the recombinant protein (bottom). Note the hH_v_1 truncated product produced when the amber stop codon is replacing F247 in the absence of the aminoacyl-tRNA synthetase/tRNA plasmid and Acd.

## Results

### We generated a library of hH_v_1 constructs with Acd incorporated across the different structural domains

Because available H_v_1 structural models were produced from either truncated or modified proteins (11, 12), we used an AlphaFold model of the full-length, dimeric hH_v_1 to guide Acd site selection (28, 29) (Figure 1B and Dataset S1). The model recapitulates the expected hH_v_1 architecture with high confidence: S0 as a short helix parallel to the membrane, S1-S4 transmembrane segments forming a canonical VSD, and a C-terminal helix extending from S4 that forms the CC intersubunit interface. In addition, two helices in the N-terminal domain (P1 and P2) were predicted (Figure 1C), albeit with low confidence. The dimer interface in the membrane is primarily formed by S4-S4 contacts (Figure S1A), although S1-S1 contacts have been observed experimentally (9, 13). This model appears to represent an intermediate state, with the first S4 positive charged residue R208 proximal to D112 in the selectivity filter, and the third S4 positive charged residue R211 proximal to F150 in the charge-transfer center (Figure S1B). Although the model’s accuracy and functional state are unknown, we used it as a rough starting point for selecting positions to incorporate Acd.

To test the feasibility and specificity of Acd incorporation in hH_v_1, we initially selected two positions in the intracellular domains: A18 in the N-terminal domain and F247 in the CC. For each position, the wild-type codon was replaced with an amber stop codon (TAG). Bacteria co-transformed with the Acd RS/tRNA pair and hH_v_1-TAG constructs expressed fluorescent full-length protein only when Acd was present in the culture medium (Figure 1D). We next generated a library of 14 constructs in which Acd was incorporated into every structural domain: the N-terminal domain, each helix of the VSD, and the CC (Figure 2A-B). Remarkably, most of the 14 hH_v_1-Acd proteins expressed at levels comparable to the control (No TAG) (Figure 2C and S2). As expected, several hH_v_1-Acd proteins also produced TAG-truncated products (Figure 2C). Increasing the Acd concentration in the *E. coli* growth medium did not reduce the fraction of truncated protein (Figure S3). One of the constructs, hH_v_1-C107Acd, showed a weak in-gel fluorescence signal, although the Western blot signal was comparable to that of other constructs.

**Figure 2.**
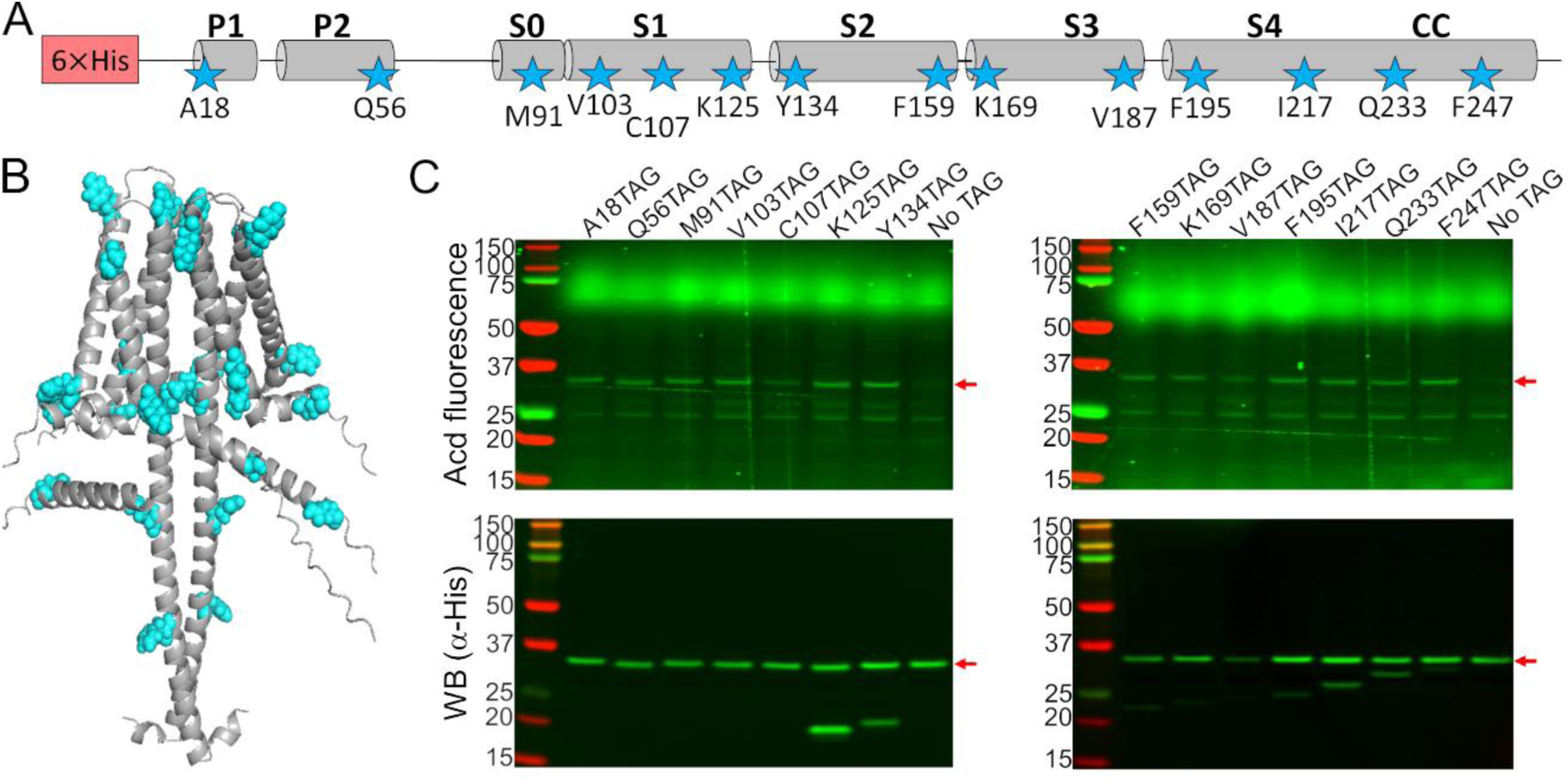
Acd was incorporated into 14 positions in the hH_v_1 sequence. (A) Cartoon showing the amino acids selected to be replaced by an amber stop codon in the hH_v_1 secondary structure to incorporate Acd (stars). (B) AlphaFold dimer hH_v_1 structural model with the amino acids selected to be replaced by Acd as cyan spheres. (C) Acd fluorescence gel (top) and Western blot (bottom) of cellular extracts after separation by SDS-PAGE showing the expression of the Acd-labeled hH_v_1 mutants (red arrows). The shorter detected bands in the WB for some constructs are produced by translation termination. The No TAG lane contains an extract from cells grown under identical conditions (in the presence of Acd and the aminoacyl-tRNA synthetase/tRNA pair) and expressing hH_v_1 without an amber stop codon.

### We successfully purified stable and functional hH_v_1 labeled with Acd at 12 of the 14 selected positions

We attempted to purify all expressed hH_v_1-Acd constructs by immobilized metal affinity chromatography using the detergent Anzergent 3-12 (20, 21). For 12 of 14 constructs, purification yielded a single fluorescent band on SDS-PAGE (Figure 3A and S4). In contrast, no purified protein was detected for the V187Acd or Q233Acd constructs (Figure S4). Because hH_v_1-Q233Acd was robustly expressed (Figure 2C), its loss during purification suggests reduced stability. The hH_v_1-Acd yields were construct-dependent (Figure 3B), and the TAG-truncated products did not co-purify with the full-length protein (Figure 3A and S4).

**Figure 3.**
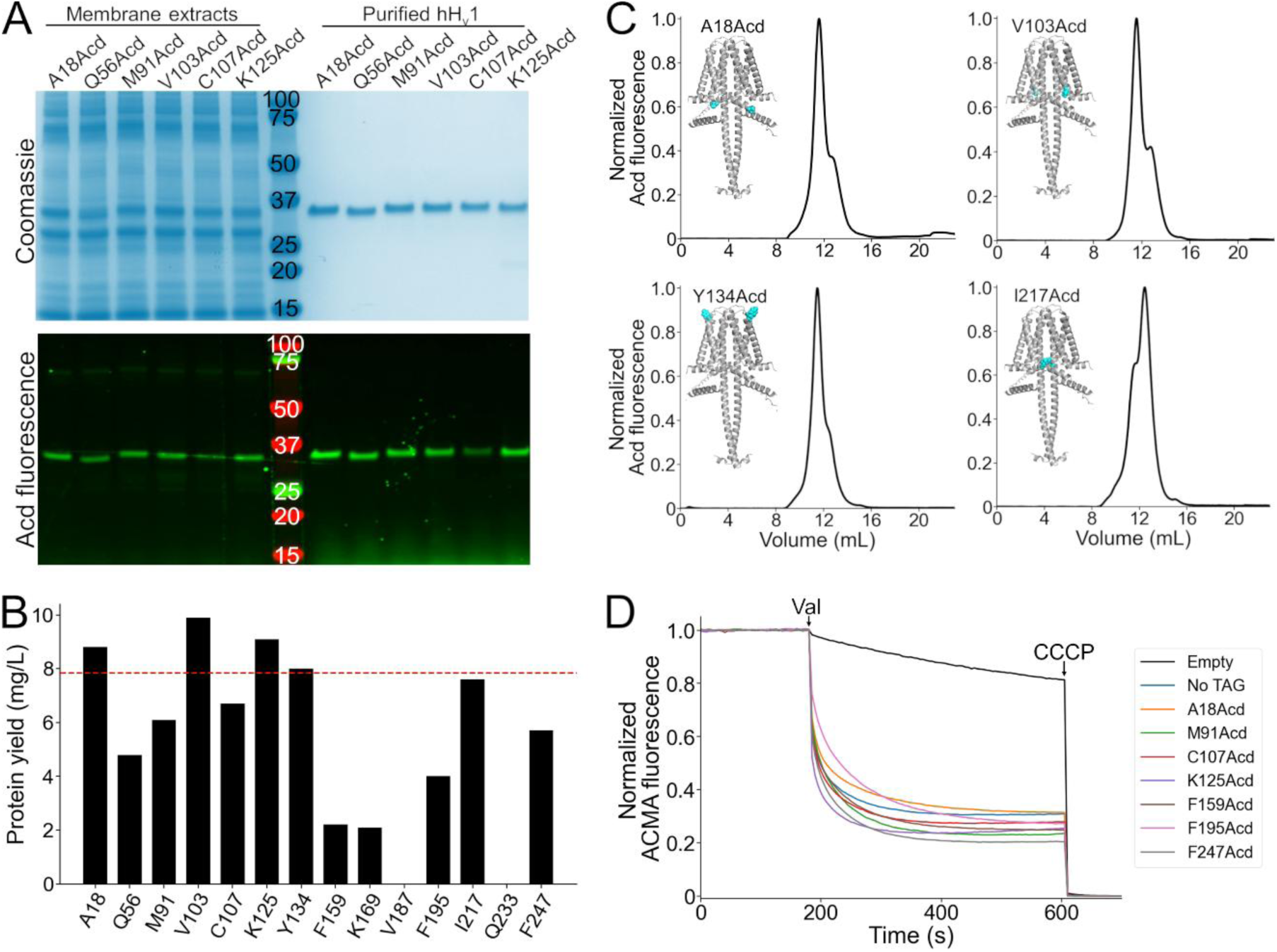
Purification and functional measurements of hH_v_1-Acd proteins. (A) Coomassie-stained (top) and Acd fluorescence (bottom) gels after separation by SDS-PAGE showing representative samples of the membrane extracts after solubilization (left) and the final purified protein after immobilized metal affinity chromatography (right). (B) Protein yield of the hH_v_1 protein with Acd replacing the amino acid at the indicated position. Note that V187Acd and Q233Acd contained no detectable protein after purification. The red line indicates the protein yield obtained from hH_v_1 without an amber stop codon in the presence of Acd and the aminoacyl-tRNA synthetase/tRNA pair (No TAG). (C) Representative fluorescence-detection size-exclusion chromatograms of the purified hH_v_1 proteins with Acd incorporated at the indicated amino acid position. Insets: AlphaFold dimer models, with the amino acid replaced by Acd shown as cyan spheres. (D) Representative liposome proton flux assay of asolectin proteoliposomes containing the indicated hH_v_1 protein. All purified proteins produced ACMA fluorescence quenching after the addition of valinomycin (Val). The protonophore CCCP was added at the end of the experiment as a control. The No TAG sample corresponds to proteoliposomes containing hH_v_1 without an amber stop codon expressed in the presence of Acd and the aminoacyl-tRNA synthetase/tRNA pair.

We then evaluated the stability and homogeneity of the hH_v_1-Acd proteins using fluorescence-detection size exclusion chromatography (FSEC) (30). Similar to the control (Figure S5), most hH_v_1-Acd proteins showed a predominant main peak (∼11.5 mL), which corresponds to the dimer (Figure S5), followed by a minor peak at higher elution volumes (Figure 3C; Table S1). The proportion of these two peaks varied between constructs (Figure S6). To determine whether the purified hH_v_1-Acd proteins were functional, we reconstituted the FSEC fractions containing both peaks in asolectin liposomes and assessed their function using a liposome proton flux assay (20, 31). All 12 reconstituted constructs produced ACMA fluorescence quenching upon valinomycin addition, indicating that our purified proteins formed functional proton-permeable channels (Figure 3D and S7). These results highlight the versatility of GCE for incorporating Acd as a fluorescent probe for structural studies of hH_v_1.

### Acd was relatively insensitive to the hydrophobicity of its local environment

To explore how the local environment can alter Acd’s spectral properties, we measured the emission spectra of the free amino acid in different solvents (Figure 4A and Table S2). The environmental sensitivity of Acd has been studied previously (24, 27, 32), but we expanded this work here by using the aprotic solvent ethyl acetate (EtAc) and a series of alcohols with varying alkyl chain lengths. Consistent with a general solvent effect produced by polarity (16), the emission spectrum was blue-shifted by approximately 20 nm in EtAc, the least polar solvent measured, relative to water (Figure 4A and Table S2). Alcohols produced a consistently smaller blue shift, independent of their alkyl-chain length (Figure 4A). The spectrum in Buffer-H2, our standard experimental solution containing detergent micelles, was only very slightly blue-shifted relative to water (Figure 4A), suggesting that Acd did not substantially partition into the hydrophobic core of the detergent micelles. We also tested whether solvent polarity affects the fluorescence lifetime of Acd using time-correlated single photon counting (TCSPC). The fluorescence lifetime of Acd showed a similar trend to the spectral shifts (Figure 4B): EtAc shortened the lifetime markedly, Acd exhibited shorter lifetimes in alcohols than in water, and Acd in Buffer-H2 had a lifetime similar to that measured in water.

**Figure 4.**
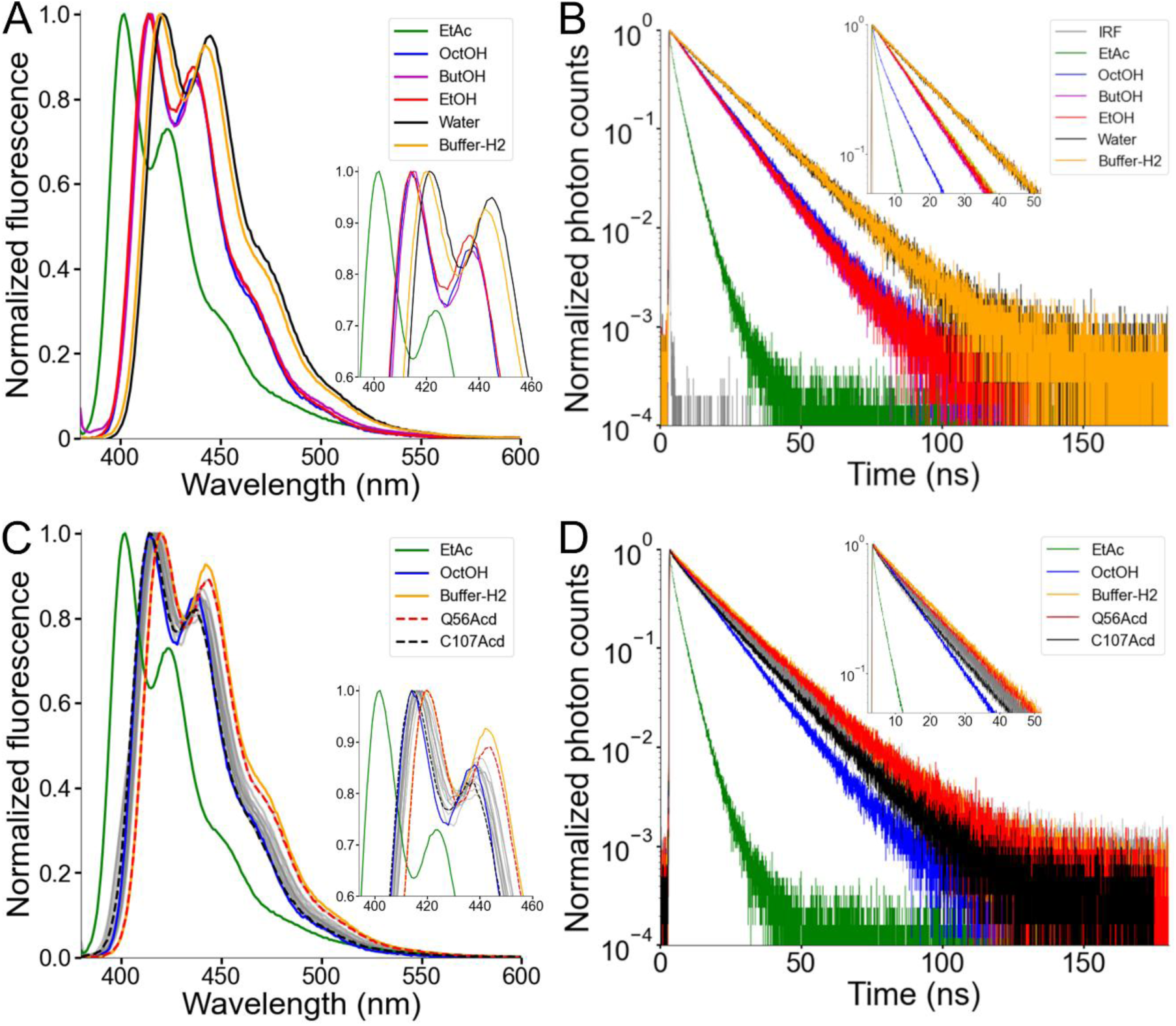
Acd showed a low environmental sensitivity in hH_v_1. (A) Normalized fluorescence emission spectra of the Acd amino acid dissolved in the indicated solvent (Excitation = 370 nm). EtAc: ethyl acetate, OctOH: 1-octanol, ButOH: 1-butanol, EtOH: ethanol. (B) Normalized fluorescence decay of the Acd amino acid dissolved in the indicated solvent. (C) Normalized fluorescence emission spectra of the hH_v_1-Acd proteins in Buffer-H2 (grey). The Acd amino acid spectra in selected solvents are also shown for reference. Q56Acd and C107Acd spectra are shown with broken lines. The emission maxima are listed in Table S3. (D) Normalized fluorescence decay of the hH_v_1-Acd proteins in Buffer-H2 (grey). The Acd amino acid decays in selected solvents are also shown for reference. Q56Acd and C107Acd decays are shown in red and black, respectively.

The spectral blue-shift and short lifetime of Acd in EtAc suggest that Acd may be able to report on the local environment at different sites in hH_v_1, especially those that are surface-exposed versus those that face the lipid core. The emission spectra of the hH_v_1-Acd proteins showed small, site-specific shifts (Figure 4C and Table S3). The local environment of Acd in hH_v_1 spanned the range between those observed for free Acd in Buffer-H2 and in alcohols, with Q56Acd in the N-terminal intracellular domain and C107Acd in S1 showing the most red-shifted and blue-shifted spectra, respectively (Figure 4C). Consistent with the spectra, the fluorescence lifetimes of the hH_v_1-Acd proteins fell between those measured for free Acd in Buffer-H2 and in alcohols, with Q56Acd exhibiting the slowest, and C107Acd the fastest, lifetimes (Figure 4D). While small, the differences in spectral properties of Acd in hH_v_1 agree with their predicted location in the structural model, with C107 being the site closest to the detergent micelles’ hydrophobic core. The low sensitivity to polarity and pH changes (33) of Acd in different hH_v_1 local environments makes it well suited for measuring FRET (27, 34).

### FRET between Trp/Tyr and Acd in hH_v_1 suggested that the proteins are properly folded

Trp and Tyr have fluorescence emission spectra that overlap with the Acd absorption spectrum and therefore can act as FRET donors to Acd, with relatively short *R_0_* values (23.5 Å for Trp; 20.9 Å for Tyr) (22) (Figure 5A). Each hH_v_1 monomer contains four Trp and four Tyr residues (yellow spheres in Figure 5B). In this structural model, the Trp residues are located in the N-terminal domain (W4, W38, and W45) and midway along transmembrane segment S4 in the VSD (W207). The Tyr residues are located in the N-terminal domain (Y35 and Y42) and near the extracellular end of transmembrane segment S2 (Y134 and Y141).

**Figure 5.**
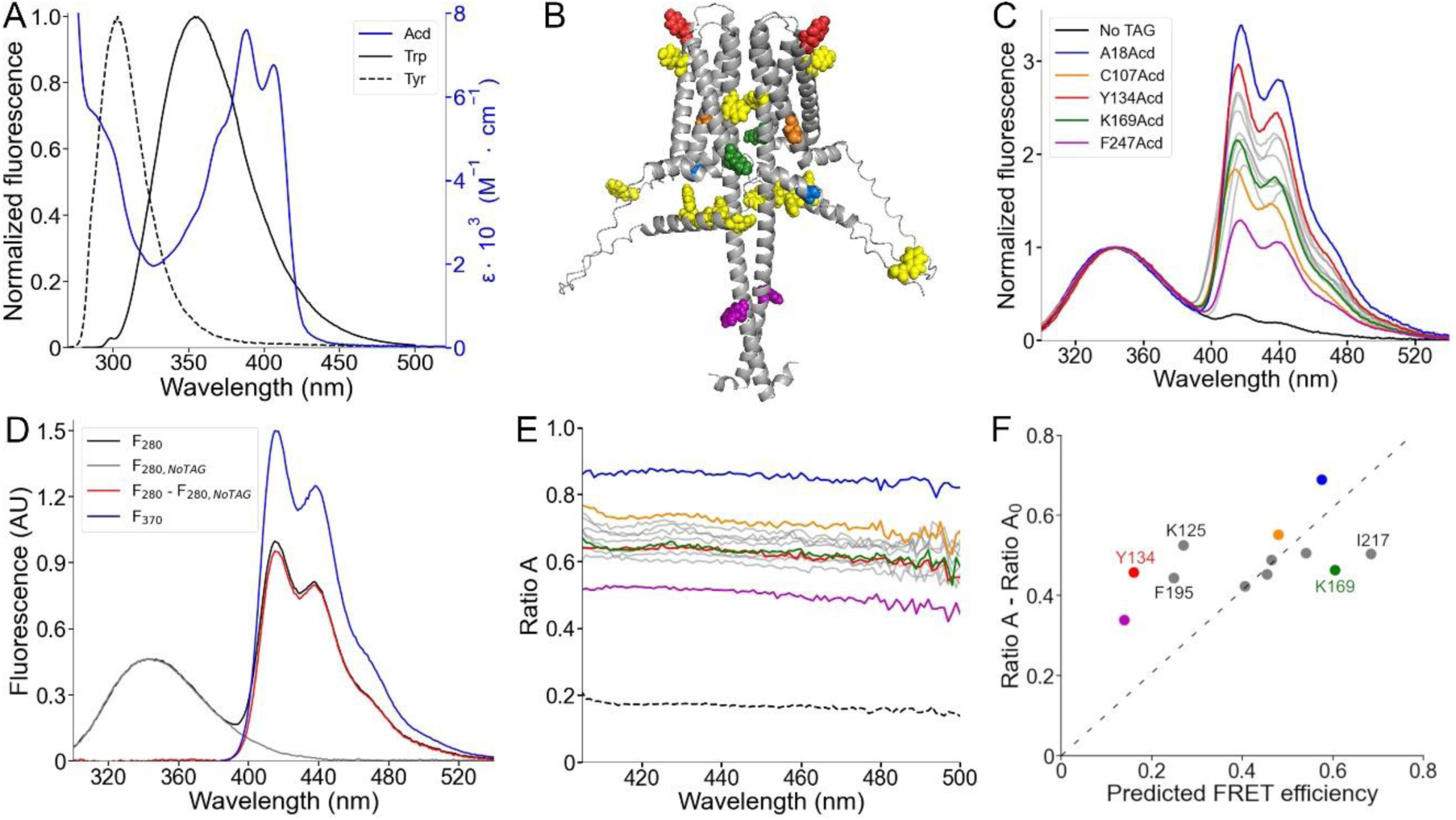
Spectral FRET analysis using Trp and Tyr donors and Acd as acceptor. (A) Fluorescence emission spectra of the intrinsic fluorescent amino acids Trp (black) and Tyr (black, broken lines), along with the Acd absorption spectrum in Buffer-H2 (blue). (B) AlphaFold dimer hH_v_1 structural model with the amino acids colored according to (C) and the native tryptophan and tyrosine residues shown as yellow spheres. (C) Normalized fluorescence spectra of the purified hH_v_1-Acd proteins (Excitation = 280 nm). The spectra were normalized by the Trp/Tyr fluorescence. The colors correspond to the indicated hH_v_1-Acd protein, and the remaining spectra are shown as grey traces in the figure. The No TAG sample (black) contains purified hH_v_1 protein without any amber stop codon expressed in the presence of Acd and the aminoacyl-tRNA synthetase/tRNA pair. (D) Example of the spectral FRET analysis procedure. The fluorescence spectrum of hH_v_1-Y134Acd obtained when exciting at 280 nm (black, *F_280_*) minus the normalized spectrum at the same excitation wavelength of the No TAG sample (grey, *F_280,NoTAG_*) produced the Acd emission spectrum shown in red (*F_280_ – F_280,NoTAG_*). *Ratio A* values were calculated by dividing this spectrum by the hH_v_1-Y134Acd fluorescence spectrum obtained by direct excitation at 370 nm (*F_370_*). (E) *Ratio A* traces of the hH_v_1-Acd proteins colored according to (C). The ratio between the spectra of free Acd amino acid excited at 280 and 370 nm (*Ratio A_0_*) is shown as broken lines. (F) Mean *Ratio A* – *Ratio A_0_* values (410-480 nm; N=4) as a function of the predicted FRET efficiencies from the AlphaFold hH_v_1 structural model, colored according to (C). The broken line is the best fit to the equation (*Ratio A* – *Ratio A_0_*) = m(Predicted FRET efficiency).

To quantify Trp/Tyr to Acd FRET, we used spectral FRET analysis (35). The total emission spectrum from Trp/Tyr and Acd was collected (Figure 5C). The Acd emission spectrum was extracted by subtracting a scaled Trp/Tyr spectrum collected from control wild-type protein (*F_280,NoTAG_*) (Figure 5D and S8). For each Acd-containing construct, the ratio of the extracted spectrum (*F_280_ – F_280,NoTAG_*) to the Acd spectrum with direct excitation (*F_370_*) was calculated as *Ratio A* (Figure 5D and 5E). Because *Ratio A* is not wavelength-dependent, it reports the linearity of the detectors and the absence of significant contamination from other sources of fluorescence. The *Ratio A* component caused by the direct excitation of Acd (termed *Ratio A_0_*) was measured with free Acd (36) (Figure 5E and S9). The difference between *Ratio A* and *Ratio A_0_* is directly proportional to the FRET efficiency.

As expected, the *Ratio A* values for our hH_v_1-Acd proteins were largely flat across emission wavelengths for our hH_v_1-Acd proteins (Figure 5E). For all Acd sites, *Ratio A* was greater than *Ratio A_0_* (dashed line in Figure 5E), reflecting FRET between Trp/Tyr and Acd at every site in hH_v_1. Most *Ratio A* values were distributed between 0.60 and 0.80, with two clear exceptions: A18Acd (blue) and F247Acd (magenta) (Figure 5E). A18Acd showed the highest *Ratio A*, consistent with the multiple Trp/Tyr residues in the N-terminal domain. Conversely, F247Acd showed the lowest *Ratio A*, consistent with its location in the CC, far from most Trp/Tyr residues (Figure 5B).

We next compared *Ratio A – Ratio A_0_* values with FRET efficiencies predicted by the AlphaFold structural model. We modeled rotamer ensembles for Trp, Tyr, and our Acd sites in the hH_v_1 structural model using chiLife (37) and calculated donor-acceptor distance distributions. These distributions were then used to calculate FRET efficiencies (details in SI Text). The experimentally determined *Ratio A – Ratio A_0_* values showed a moderate correlation (Pearson’s r = 0.48) with the calculated efficiencies (Figure 5F). Using the predicted intrasubunit FRET efficiencies reduced the Pearson’s r value to 0.18 (Figure S10), suggesting that the measured FRET contains information about subunit organization.

There were two spatially clustered deviations between experiment and model. Positions at the extracellular ends of the VSD helices (Y134 in S2, K125 in S1, and F195 in S4) showed higher experimental FRET than predicted, indicating shorter effective donor-acceptor distances relative to the model. Conversely, positions on the intracellular side of the VSD (K169 in S3 and I217 in S4) showed lower experimental FRET than predicted, implying longer effective donor-acceptor distances. Together, the spatial clustering of residues with similar deviations indicates specific deficits in the AlphaFold model or the failure to quantify the protein dynamics.

### Zinc binding changed the conformation of hH_v_1

We next asked whether FRET from Trp/Tyr to Acd could report conformational changes upon addition of Zn^2+^, the classical inhibitor of H_v_1 proton currents in cells (7, 38, 39). The two experimental structures of H_v_1 were resolved with this cation bound to the extracellular side of the VSD (11, 12). We observed site-specific spectral changes in the hH_v_1-Acd proteins in the presence of Zn^2+^ (Figure 6 and S11).

**Figure 6.**
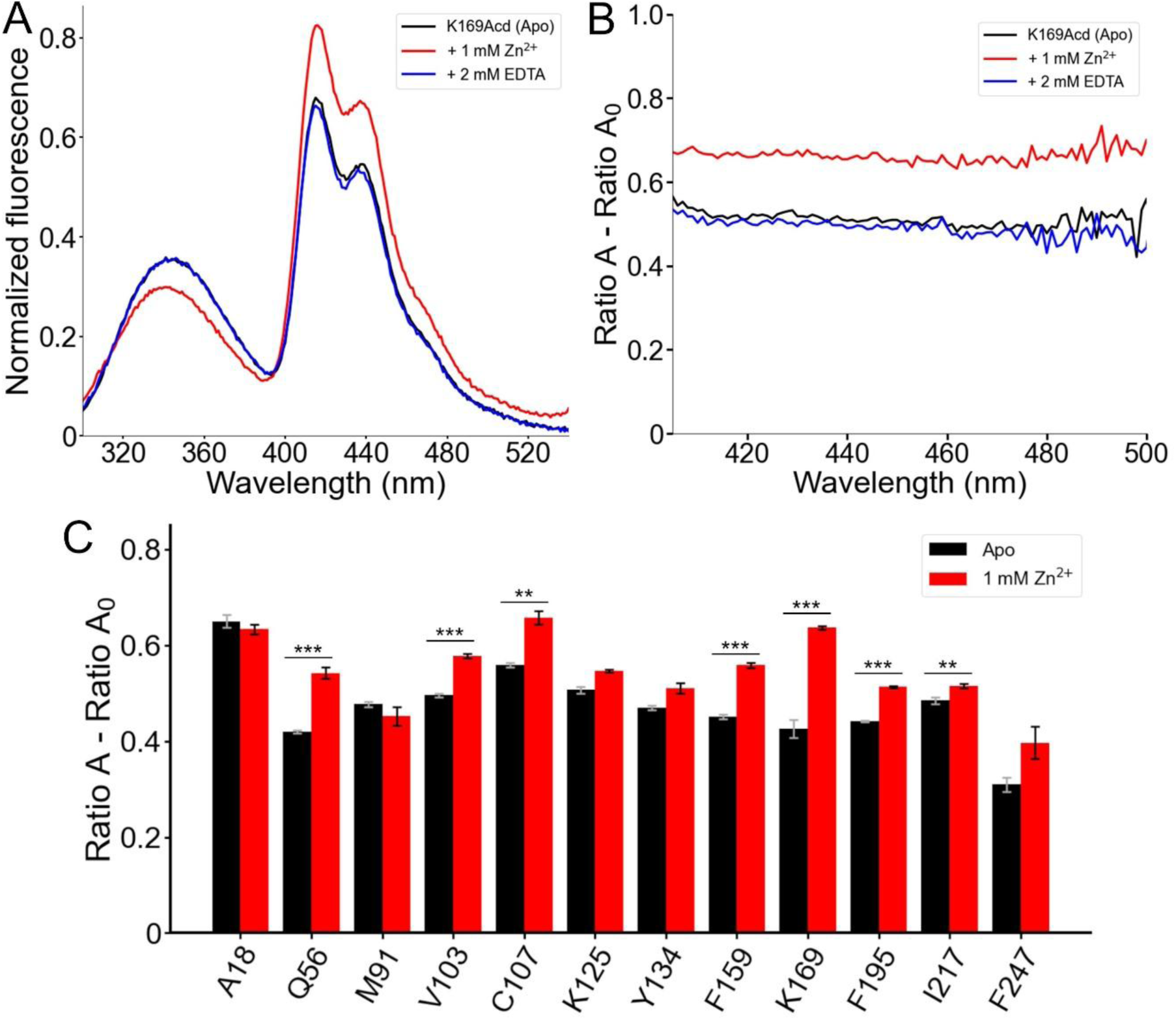
FRET between Trp/Tyr and Acd reports a conformational rearrangement in response to Zn^2+^. (A) Normalized fluorescence emission spectra of hH_v_1-K169Acd (Excitation = 280 nm) in Buffer-H2 in the absence of Zn^2+^ (black, Apo), in the presence of 1 mM Zn^2+^ (red), or in the presence of 1 mM Zn^2+^ and 2 mM EDTA (blue). Spectra were normalized by the maximum intensity of the Acd emission spectrum of the same sample excited at 370 nm. (B) Spectral FRET analysis of hH_v_1-K169Acd in Buffer-H2 in the absence of Zn^2+^ (black, Apo), in the presence of 1 mM Zn^2+^ (red), or in the presence of 1 mM Zn^2+^ and 2 mM EDTA (blue). (C) Spectral FRET analysis of the hH_v_1-Acd proteins in Buffer-H2 in the absence (black, Apo) or presence of 1 mM Zn^2+^ (red). N=4, ***p<0.005, **p<0.01.

The largest changes occurred for hH_v_1-K169Acd, where 1 mM Zn^2+^ caused a decrease in the Trp/Tyr emission and an increase in the Acd emission (Figure 6A). This decrease in donor fluorescence, paired with increased acceptor fluorescence, is the hallmark of increased FRET and was corroborated by spectral analysis (Figure 6B). Additionally, a slight blue shift of the Trp/Tyr fluorescence was observed for all hH_v_1-Acd proteins in the presence of Zn^2+^ (Figure S11). These changes were completely reversed by the addition of EDTA, confirming that the changes arise from a reversible conformational change upon Zn^2+^ binding (Figure 6A and 6B). Large changes in effective FRET efficiency were observed with Acd incorporated at Q56 in the N-terminal domain and C107, F159, and K169 in the VSD (Figure 6C). Notably, these positions are intracellular, even though H_v_1 currents are inhibited by extracellular Zn^2+^ (7, 40). These results are consistent with previous studies showing that some extracellular H_v_1 inhibitors, such as Zn^2+^ or AGAP, induce long-range structural changes that propagate to the channel’s intracellular vestibule (41, 42) and persist in detergent micelles (12).

## Discussion

Incorporating Acd into hH_v_1 by GCE opens new possibilities for studying the structure and function of this channel. We systematically incorporated Acd at multiple positions in hH_v_1 and confirmed protein stability and function after purification. 12 of 14 hH_v_1-Acd proteins produced acceptable yields upon purification and were functional, despite being a human membrane protein expressed in *E. coli*. Two factors likely underlie the robustness of our system: (i) an optimized purification protocol for hH_v_1 (20, 21) that offsets the expected reduction in yield associated with GCE (22, 25), and (ii) the high efficiency and specificity of the evolved aminoacyl-tRNA synthetase for incorporating Acd (17, 27).

Fluorescence spectroscopy has been used to detect conformational changes in H_v_1. Sites at the extracellular portion of S4 (43) and S1 (44) in the VSD of *Ciona intestinalis* H_v_1 have been labeled with the fluorophores Alexa488 and tetramethylrhodamine (TAMRA) through cysteine-selective reagents to capture conformational changes during activation using voltage-clamp fluorometry. Similarly, the fluorescent noncanonical amino acid 3-(6-acetylnaphthalen-2-ylamino)-2-aminopropanoic acid (Anap) has been incorporated at several sites in the S4 in hH_v_1 (45) to study conformational changes in response to voltage and pH. In these cases, the fluorophore’s environmental sensitivity has been used as a readout of conformational changes. By contrast, FRET provides richer structural information due to its distance dependence. Single-molecule FRET has been measured in purified hH_v_1 labeled with Cy3 and Cy5 to track the VSD movements in response to voltage, pH, cholesterol, and zinc (14, 46). The small size and minimal linker of Acd, along with its low environmental sensitivity and high photostability, are advantageous for measuring FRET in hH_v_1 compared with previously used fluorophores.

The high number of FRET donors, 4 tryptophans and 4 tyrosines per subunit, limits the utility of the Trp/Tyr-Acd FRET pair as a strategy to measure conformational rearrangements in hH_v_1. Our library of a dozen single Acd-incorporating constructs, with sites distributed across the entire primary sequence of the protein, opens the door to using Acd as a FRET donor. We have previously used time-resolved transition metal ion FRET (tmFRET), which utilizes a transition metal ion as a FRET acceptor (27, 34), to measure conformational energetics in the bacterial cyclic nucleotide-gated ion channel SthK (47). However, in that work, we incorporated Acd at a single position in the intracellular C-terminal domain of SthK. With Acd incorporated at multiple sites in full-length hH_v_1, it will be possible to interrogate conformational changes across the protein’s different structural domains using Acd as a tmFRET donor to understand its molecular mechanisms.

## Materials and Methods

### Protein expression and purification

hH_v_1 was expressed and purified according to the published protocol with minor modifications (20, 21). Briefly, fresh competent BL21-Gold(DE3) cells (Agilent) were co-transformed with the His-EK-hH_v_1-C107A-C249A.pET15-b (hH_v_1-Cysless, referred to as No TAG) and MjA9Acd-RS.pDule2 (22, 23) plasmids (sequences in SI Text) using the heat shock method. For expressing hH_v_1-Acd proteins, the codon corresponding to the selected position was changed to the amber stop codon (TAG) by site-directed mutagenesis. All plasmids were sequenced before use. Pre-cultures were grown overnight in LB medium supplemented with 0.2% glucose, 0.4 mg/mL ampicillin, and 0.1 mg/mL spectinomycin at 37 °C and 250 rpm. The next day, cultures were diluted 1:100 in complete autoinduction medium (48) supplemented with 0.4 mg/mL ampicillin and 0.1 mg/mL spectinomycin, and then grown at 37 °C and 250 rpm until an OD_600_ of 1.5 was reached. The protein expression level was proportional to the Acd concentration in the culture medium (Figure S12). We chose 0.6 mM Acd in the culture medium as optimal for hH_v_1 expression, as higher concentrations only slightly increased the final biomass (Figure S1B). Cultures supplemented with 0.6 mM Acd were grown overnight at 20 °C and 250 rpm. Cells were then harvested, resuspended in Buffer-H1 (50 mM Tris, 150 mM NaCl, 1 mM benzamidine, 0.17 mg/mL PMSF, pH 8.0) supplemented with 0.5 mg/mL lysozyme, and incubated for 30 min at 4 °C before being flash-frozen. After thawing, the extracts were supplemented with 5 mM MgCl_2_, fresh protease inhibitors, and 12.5 μg/mL DNase I, incubated for 1 h at 4°C, sonicated, and centrifuged at 100,000g at 4 °C for 1 h. The membrane pellet was resuspended in Buffer-H1 and stored at -80 °C until further use. After thawing, the membranes were solubilized with 1.5% Anzergent 3-12 (Anatrace) (Anz3-12) at room temperature for 1 hour. Insoluble material was removed by centrifuging at 100,000g and 4 °C for 1 h, and the solubilized membrane extract was incubated with His-Pur Ni-NTA (Thermo Scientific) resin equilibrated with Buffer-H2 (50 mM Tris, 150 mM NaCl, 12 mM Anz3-12, pH 8.0) for 1 hour at room temperature. Resin was collected in a gravity column and washed with 20 column volumes (CV) of Buffer-H2, followed by 16 CV of Buffer-H2 with 90 mM imidazole. The hH_v_1 protein was eluted with 20 CV of Buffer-H2 with 0.5 M imidazole, was concentrated with 50kDa cut-off centrifugal filters, and imidazole was removed by FSEC (Ex/Em = 385/450 nm) in an ENrich SEC 650 (Bio-Rad) column with Buffer-H2. The hH_v_1-containing fractions were concentrated to 0.5-1.5 mg/mL, depending on the amount of purified protein, and then aliquoted, flash-frozen, and stored at -80 °C until use.

### Reconstitution and fluorescence proton flux assays

The hH_v_1-Acd samples were reconstituted and assayed using a liposome proton flux assay, as described previously, with minor modifications (20). Briefly, soy polar lipid extract (Avati) in chloroform was dried overnight at room temperature in a rotary evaporator under vacuum and then hydrated in Buffer-K (20 mM HEPES, 150 mM KCl, and 1 mM EDTA, pH 7.0) to a concentration of 10 mg/mL. The vesicles were sonicated in a water bath until the solution became translucent, aliquoted, flash-frozen, and stored at -80 °C until further use. Reconstitutions were performed at a 1:100 (protein:lipid, by mass) ratio by mixing 12.5 μg of purified hH_v_1-Acd with 1.25 mg of liposomes in a final volume of 330 μL Buffer-K containing 8 mM Anz3-12. The mixture was incubated at room temperature for 1 h, then diluted with 5 mL of Buffer-K and incubated for 30 min. The mixture was further incubated with three cumulative pulses of 100 mg Bio-Beads SM2 (Bio-Rad), each for 1 h at room temperature. After the final Bio-Beads pulse, the mixture was incubated overnight at 4 °C. The next day, two successive 100 mg additions of Bio-Beads were performed, with a 1-hour incubation at room temperature following each addition. Bio-Beads were removed in a gravity column, the mixture was diluted with 10 mL of Buffer-K, and then centrifuged at 150,000g for 2 hours at 4 °C. The proteoliposome pellets were resuspended in 250 μL of Buffer-K, aliquoted, flash-frozen, and stored at -80 °C until further use. On the day of the liposome proton flux assays, samples were thawed at 37 °C and then placed on ice. The sample (40 μL) was diluted in Buffer-Na (20 mM HEPES, 150 mM NaCl, 1 mM EDTA, pH 7.0). 9-Amino-6-chloro-2-methoxyacridine (ACMA; Sigma) was added to a final concentration of 2 μM from a 2 mM stock solution in DMSO, and the mixture was incubated at room temperature for 5 minutes before the fluorescence measurement began. The baseline was recorded for 3 min (*F_max_*). Subsequently, 10 nM of valinomycin (Cell Signaling) was added from a 10 μM stock solution in DMSO, followed by 1 μM carbonyl cyanide m-chlorophenyl hydrazone (CCCP; Sigma) from a 1 mM stock solution in DMSO to record *F_min_*. The final volume was 2 mL. The fluorescence signal (*F(t)*) was normalized according to the equation (*F(t)*-*F_min_*)/(*F_max_*-*F_min_*). Fluorescence measurements were performed at 5-second intervals with a 2-second integration time using a Fluorolog-3 spectrometer (Horiba) configured to an excitation wavelength of 410 nm (5 nm bandpass) and an emission wavelength of 490 nm (1 nm bandpass) at room temperature.

### Absorption and fluorescence spectra recordings

The absorption and fluorescence spectra were recorded in the indicated solvents or fresh Buffer-H2 at room temperature. Absorption spectra were recorded in a DU 800 Spectrometer (Beckman Coulter) with a wavelength interval of 0.5 nm and a scan speed of 600 nm/min. Fluorescence emission spectra were measured using a Fluorolog-3 spectrometer (Horiba) with an integration time of 0.1 s, 1 nm increments, and excitation and emission slits of 5 nm. For solubilizing Acd in different solvents, a saturated solution of the amino acid was prepared by dissolving 0.5 mg of Acd powder in the respective solvent. Acd spectra were corrected by subtracting a blank of the solvent. hH_v_1-Acd proteins were measured at concentrations ranging from 180 to 240 nM in Buffer-H2. Each spectrum was corrected by subtracting a blank with the solvent or Buffer-H2 before adding the protein sample to the cuvette.

### Spectral FRET analysis

We used our previously established spectral FRET analysis to remove contamination caused by direct excitation of Acd by 280 nm light (36). This method had the added benefit of eliminating errors arising from the recording system’s transfer function, variations in the acceptor’s quantum yield, or variations in the total concentration of fluorescent molecules. A Trp/Tyr spectrum was collected from control protein without any TAG codons (no Acd incorporation). This was used to subtract the Trp/Tyr emission spectra collected at 280 nm for each Acd-incorporating protein. This yielded the extracted Acd spectrum, *F_280_*, that had two components: the component due to direct excitation of Acd, 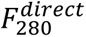, and the component due to FRET 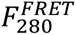. *F_280_* was normalized by the total Acd emission with 370 nm excitation, *F_370_*. The resulting ratio, termed *Ratio A*, can be expressed as: 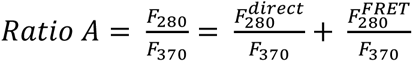. The direct excitation component, 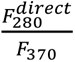, termed *Ratio A_0_*, was measured using free Acd amino acid in Buffer-H2. We then quantified the relative FRET efficiency as 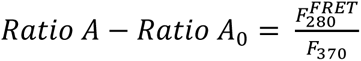. This quantity is directly proportional to FRET efficiency. An average *Ratio A* value was calculated between 410 and 480 nm.

### Time-resolved fluorescence measurements

TCSPC measurements were performed in a FluoTime 300 spectrometer equipped with a PMA Hybrid 40 detector and LDH-P-C-375 laser diode head (PicoQuant) at room temperature. Emitted photons were detected at 446 nm with the emission polarizer at the magic angle. The TCSPC measurements were performed in the indicated solvent or fresh Buffer-H2. The hH_v_1-Acd samples were measured at a concentration of 1 μM and the Acd amino acid at 500 nM. The count rate was always less than 1% of the excitation repetition rate to avoid pile-up distortions of the fluorescence decay. The IRF was measured using a dilute Ludox solution in water.

## Acknowledgments

We thank the Oregon State University GCE4ALL (Center for Genetic Code Expansion for All) for their long-standing collaboration. We also appreciate the excellent technical support provided by Dr. James Petersson and Kyle D. Shaffer (University of Pennsylvania) with the Acd synthesis. This work was supported by the National Institutes of Health under award numbers R01EY037223 and R35GM145225 to S.E.G., and R01EY010329 and R35GM148137 to W.N.Z. E.M.C. is a Pew Latin American Fellow.

## Supplementary Text

### Calculation of the FRET efficiency from the structural model

We calculated the FRET efficiency using the AlphaFold model of the full-length hH_v_1 (Dataset S1). For each Acd position, the measured steady-state FRET would correspond to the mean FRET efficiency (*E_FRET_*), which will be proportional to the *Ratio A – Ratio A_0_* value obtained from the spectral FRET analysis.

For *n* number of donors and two acceptors in the hH_v_1 dimer:

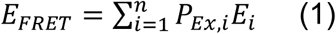

where *P_Ex,i_* is the probability of excitation and *E_i_* is the FRET efficiency of the *i*th donor. *P_Ex,i_* is the number of photons absorbed by the *i*th donor divided by the total number of photons absorbed by the total number *n* of donors, which is quantified by the donor’s extinction coefficient. Supposing the same extinction coefficient for each of the Trp (*ε_W_* = 5,501 M^-1^ cm^-1^) and Tyr (*ε_Y_* = 1,209 M^-1^ cm^-1^) (Supplementary Reference 1), equation 1 can be written as:

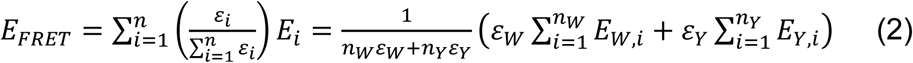

where the sum of the total number of Trp (*n_W_*) and Tyr (*n_Y_*) is equal to *n*. The efficiency of FRET for the *i*th donor can be expressed as the sum of rates of energy transfer to each of the *j* = 2 Acd acceptors, *k_ij_*, divided by the total rates of photon emission. Therefore, the right terms of equation 2 can be expressed as:

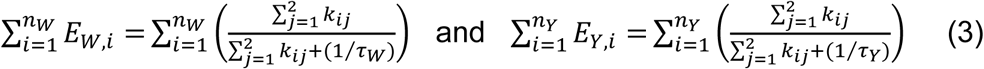

where the Trp lifetime (*τ_W_* = 3.1 ns) and tyrosine lifetime (*τ_Y_* = 3.6 ns) in the absence of donors are considered constant (Supplementary Reference 1). Finally, each energy transfer rate *k_ij_* is a function of the distance *r_ij_* between the *i*th donor and the *j*th acceptor:

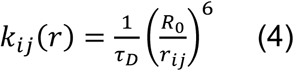

where the donor lifetime τ*_D_* and *R_0_* would be *τ_W_* and 23.5 Å or *τ_Y_* and 20.9 Å for Trp or Tyr residues, respectively. For determining the *R_0_* values from the spectra overlap, we used an orientation factor value of 2/3, a refraction index of 1.3346 measured for BufferH2 using a refractometer, and quantum yields of 0.12 and 0.13 for Trp and Tyr, respectively (Supplementary Reference 1). Calculating *E_FRET_* using the above equations considers a fixed distance between donors and acceptors. A better approach is to consider the distance distributions *P_ij_(r)* produced by the different rotameric states of the FRET pair, which can be modeled easily using chiLife (Supplementary Reference 2). Then, equation 4 will be replaced by equation 5:

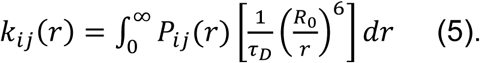

### DNA plasmid sequences in FASTA format

>His-EK-hHv1-C107A-C249A.pET15-b

TTGAGATCCTTTTTTTCTGCGCGTAATCTGCTGCTTGCAAACAAAAAAACCACCGCTACCAGCGGTGGTTTGTTTGCCGGATCAAGAGCTACCAACTCTTTTTCCGAAGGTAACTGGCTTCAGCAGAGCGCAGATACCAAATACTGTCCTTCTAGTGTAGCCGTAGTTAGGCCACCACTTCAAGAACTCTGTAGCACCGCCTACATACCTCGCTCTGCTAATCCTGTTACCAGTGGCTGCTGCCAGTGGCGATAAGTCGTGTCTTACCGGGTTGGACTCAAGACGATAGTTACCGGATAAGGCGCAGCGGTCGGGCTGAACGGGGGGTTCGTGCACACAGCCCAGCTTGGAGCGAACGACCTACACCGAACTGAGATACCTACAGCGTGAGCTATGAGAAAGCGCCACGCTTCCCGAAGGGAGAAAGGCGGACAGGTATCCGGTAAGCGGCAGGGTCGGAACAGGAGAGCGCACGAGGGAGCTTCCAGGGGGAAACGCCTGGTATCTTTATAGTCCTGTCGGGTTTCGCCACCTCTGACTTGAGCGTCGATTTTTGTGATGCTCGTCAGGGGGGCGGAGCCTATGGAAAAACGCCAGCAACGCGGCCTTTTTACGGTTCCTGGCCTTTTGCTGGCCTTTTGCTCACATGTTCTTTCCTGCGTTATCCCCTGATTCTGTGGATAACCGTATTACCGCCTTTGAGTGAGCTGATACCGCTCGCCGCAGCCGAACGACCGAGCGCAGCGAGTCAGTGAGCGAGGAAGCGGAAGAGCGCCTGATGCGGTATTTTCTCCTTACGCATCTGTGCGGTATTTCACACCGCATATATGGTGCACTCTCAGTACAATCTGCTCTGATGCCGCATAGTTAAGCCAGTATACACTCCGCTATCGCTACGTGACTGGGTCATGGCTGCGCCCCGACACCCGCCAACACCCGCTGACGCGCCCTGACGGGCTTGTCTGCTCCCGGCATCCGCTTACAGACAAGCTGTGACCGTCTCCGGGAGCTGCATGTGTCAGAGGTTTTCACCGTCATCACCGAAACGCGCGAGGCAGCTGCGGTAAAGCTCATCAGCGTGGTCGTGAAGCGATTCACAGATGTCTGCCTGTTCATCCGCGTCCAGCTCGTTGAGTTTCTCCAGAAGCGTTAATGTCTGGCTTCTGATAAAGCGGGCCATGTTAAGGGCGGTTTTTTCCTGTTTGGTCACTGATGCCTCCGTGTAAGGGGGATTTCTGTTCATGGGGGTAATGATACCGATGAAACGAGAGAGGATGCTCACGATACGGGTTACTGATGATGAACATGCCCGGTTACTGGAACGTTGTGAGGGTAAACAACTGGCGGTATGGATGCGGCGGGACCAGAGAAAAATCACTCAGGGTCAATGCCAGCGCTTCGTTAATACAGATGTAGGTGTTCCACAGGGTAGCCAGCAGCATCCTGCGATGCAGATCCGGAACATAATGGTGCAGGGCGCTGACTTCCGCGTTTCCAGACTTTACGAAACACGGAAACCGAAGACCATTCATGTTGTTGCTCAGGTCGCAGACGTTTTGCAGCAGCAGTCGCTTCACGTTCGCTCGCGTATCGGTGATTCATTCTGCTAACCAGTAAGGCAACCCCGCCAGCCTAGCCGGGTCCTCAACGACAGGAGCACGATCATGCGCACCCGTGGCCAGGACCCAACGCTGCCCGAGATGCGCCGCGTGCGGCTGCTGGAGATGGCGGACGCGATGGATATGTTCTGCCAAGGGTTGGTTTGCGCATTCACAGTTCTCCGCAAGAATTGATTGGCTCCAATTCTTGGAGTGGTGAATCCGTTAGCGAGGTGCCGCCGGCTTCCATTCAGGTCGAGGTGGCCCGGCTCCATGCACCGCGACGCAACGCGGGGAGGCAGACAAGGTATAGGGCGGCGCCTACAATCCATGCCAACCCGTTCCATGTGCTCGCCGAGGCGGCATAAATCGCCGTGACGATCAGCGGTCCAGTGATCGAAGTTAGGCTGGTAAGAGCCGCGAGCGATCCTTGAAGCTGTCCCTGATGGTCGTCATCTACCTGCCTGGACAGCATGGCCTGCAACGCGGGCATCCCGATGCCGCCGGAAGCGAGAAGAATCATAATGGGGAAGGCCATCCAGCCTCGCGTCGCGAACGCCAGCAAGACGTAGCCCAGCGCGTCGGCCGCCATGCCGGCGATAATGGCCTGCTTCTCGCCGAAACGTTTGGTGGCGGGACCAGTGACGAAGGCTTGAGCGAGGGCGTGCAAGATTCCGAATACCGCAAGCGACAGGCCGATCATCGTCGCGCTCCAGCGAAAGCGGTCCTCGCCGAAAATGACCCAGAGCGCTGCCGGCACCTGTCCTACGAGTTGCATGATAAAGAAGACAGTCATAAGTGCGGCGACGATAGTCATGCCCCGCGCCCACCGGAAGGAGCTGACTGGGTTGAAGGCTCTCAAGGGCATCGGTCGAGATCCCGGTGCCTAATGAGTGAGCTAACTTACATTAATTGCGTTGCGCTCACTGCCCGCTTTCCAGTCGGGAAACCTGTCGTGCCAGCTGCATTAATGAATCGGCCAACGCGCGGGGAGAGGCGGTTTGCGTATTGGGCGCCAGGGTGGTTTTTCTTTTCACCAGTGAGACGGGCAACAGCTGATTGCCCTTCACCGCCTGGCCCTGAGAGAGTTGCAGCAAGCGGTCCACGCTGGTTTGCCCCAGCAGGCGAAAATCCTGTTTGATGGTGGTTAACGGCGGGATATAACATGAGCTGTCTTCGGTATCGTCGTATCCCACTACCGAGATATCCGCACCAACGCGCAGCCCGGACTCGGTAATGGCGCGCATTGCGCCCAGCGCCATCTGATCGTTGGCAACCAGCATCGCAGTGGGAACGATGCCCTCATTCAGCATTTGCATGGTTTGTTGAAAACCGGACATGGCACTCCAGTCGCCTTCCCGTTCCGCTATCGGCTGAATTTGATTGCGAGTGAGATATTTATGCCAGCCAGCCAGACGCAGACGCGCCGAGACAGAACTTAATGGGCCCGCTAACAGCGCGATTTGCTGGTGACCCAATGCGACCAGATGCTCCACGCCCAGTCGCGTACCGTCTTCATGGGAGAAAATAATACTGTTGATGGGTGTCTGGTCAGAGACATCAAGAAATAACGCCGGAACATTAGTGCAGGCAGCTTCCACAGCAATGGCATCCTGGTCATCCAGCGGATAGTTAATGATCAGCCCACTGACGCGTTGCGCGAGAAGATTGTGCACCGCCGCTTTACAGGCTTCGACGCCGCTTCGTTCTACCATCGACACCACCACGCTGGCACCCAGTTGATCGGCGCGAGATTTAATCGCCGCGACAATTTGCGACGGCGCGTGCAGGGCCAGACTGGAGGTGGCAACGCCAATCAGCAACGACTGTTTGCCCGCCAGTTGTTGTGCCACGCGGTTGGGAATGTAATTCAGCTCCGCCATCGCCGCTTCCACTTTTTCCCGCGTTTTCGCAGAAACGTGGCTGGCCTGGTTCACCACGCGGGAAACGGTCTGATAAGAGACACCGGCATACTCTGCGACATCGTATAACGTTACTGGTTTCACATTCACCACCCTGAATTGACTCTCTTCCGGGCGCTATCATGCCATACCGCGAAAGGTTTTGCGCCATTCGATGGTGTCCGGGATCTCGACGCTCTCCCTTATGCGACTCCTGCATTAGGAAGCAGCCCAGTAGTAGGTTGAGGCCGTTGAGCACCGCCGCCGCAAGGAATGGTGCATGCAAGGAGATGGCGCCCAACAGTCCCCCGGCCACGGGGCCTGCCACCATACCCACGCCGAAACAAGCGCTCATGAGCCCGAAGTGGCGAGCCCGATCTTCCCCATCGGTGATGTCGGCGATATAGGCGCCAGCAACCGCACCTGTGGCGCCGGTGATGCCGGCCACGATGCGTCCGGCGTAGAGGATCGAGATCTCGATCCCGCGAAATTAATACGACTCACTATAGGGGAATTGTGAGCGGATAACAATTCCCCTCTAGAAATAATTTTGTTTAACTTTAAGAAGGAGATATACCATGGGCAGCAGCCATCATCATCATCATCACAGCAGCGGCGATGATGATGATAAAATGGCGACCTGGGACGAAAAGGCGGTGACCCGTCGTGCGAAAGTTGCGCCGGCGGAGCGTATGAGCAAGTTCCTGCGTCACTTTACCGTGGTTGGTGACGATTACCACGCGTGGAACATCAACTATAAGAAATGGGAGAACGAGGAAGAGGAAGAGGAAGAGGAACAGCCGCCGCCGACCCCGGTTAGCGGCGAGGAAGGCCGTGCGGCGGCGCCGGATGTGGCGCCGGCGCCGGGTCCGGCGCCGCGTGCGCCGCTGGATTTCCGTGGCATGCTGCGTAAACTGTTCAGCAGCCACCGTTTTCAAGTTATCATTATCGCGCTGGTGGTTCTGGATGCGCTGCTGGTGCTGGCGGAGCTGATCCTGGACCTGAAGATTATCCAGCCGGATAAAAACAACTACGCGGCGATGGTTTTCCACTATATGAGCATCACCATTCTGGTGTTCTTTATGATGGAAATCATCTTCAAGCTGTTCGTTTTCCGTCTGGAGTTCTTTCACCACAAATTCGAAATCCTGGACGCGGTGGTTGTGGTTGTGAGCTTTATCCTGGATATTGTGCTGCTGTTCCAGGAGCACCAATTTGAAGCGCTGGGTCTGCTGATTCTGCTGCGTCTGTGGCGTGTTGCGCGTATTATCAACGGCATTATCATTAGCGTGAAGACCCGTAGCGAGCGTCAGCTGCTGCGTCTGAAGCAGATGAACGTTCAACTGGCGGCGAAAATCCAGCACCTGGAATTTAGCGCGAGCGAGAAGGAACAAGAGATTGAACGTCTGAACAAACTGCTGCGTCAGCACGGTCTGCTGGGCGAAGTGAACTAAGGATCCGGCTGCTAACAAAGCCCGAAAGGAAGCTGAGTTGGCTGCTGCCACCGCTGAGCAATAACTAGCATAACCCCCTTGGGGCCTCTAAACGGGTCTTGAGGGGTTTTTTGCTGAAAGGAGGAACTATATCCGGATATCCCGCAAGAGGCCCGGCAGTACCGGCATAACCAAGCCTATGCCTACAGCATCCAGGGTGACGGTGCCGAGGATGACGATGAGCGCATTGTTAGATTTCATACACGGTGCCTGACTGCGTTAGCAATTTAACTGTGATAAACTACCGCATTAAAGCTTATCGATGATAAGCTGTCAAACATGAGAATTCTTGAAGACGAAAGGGCCTCGTGATACGCCTATTTTTATAGGTTAATGTCATGATAATAATGGTTTCTTAGACGTCAGGTGGCACTTTTCGGGGAAATGTGCGCGGAACCCCTATTTGTTTATTTTTCTAAATACATTCAAATATGTATCCGCTCATGAGACAATAACCCTGATAAATGCTTCAATAATATTGAAAAAGGAAGAGTATGAGTATTCAACATTTCCGTGTCGCCCTTATTCCCTTTTTTGCGGCATTTTGCCTTCCTGTTTTTGCTCACCCAGAAACGCTGGTGAAAGTAAAAGATGCTGAAGATCAGTTGGGTGCACGAGTGGGTTACATCGAACTGGATCTCAACAGCGGTAAGATCCTTGAGAGTTTTCGCCCCGAAGAACGTTTTCCAATGATGAGCACTTTTAAAGTTCTGCTATGTGGCGCGGTATTATCCCGTGTTGACGCCGGGCAAGAGCAACTCGGTCGCCGCATACACTATTCTCAGAATGACTTGGTTGAGTACTCACCAGTCACAGAAAAGCATCTTACGGATGGCATGACAGTAAGAGAATTATGCAGTGCTGCCATAACCATGAGTGATAACACTGCGGCCAACTTACTTCTGACAACGATCGGAGGACCGAAGGAGCTAACCGCTTTTTTGCACAACATGGGGGATCATGTAACTCGCCTTGATCGTTGGGAACCGGAGCTGAATGAAGCCATACCAAACGACGAGCGTGACACCACGATGCCTGCAGCAATGGCAACAACGTTGCGCAAACTATTAACTGGCGAACTACTTACTCTAGCTTCCCGGCAACAATTAATAGACTGGATGGAGGCGGATAAAGTTGCAGGACCACTTCTGCGCTCGGCCCTTCCGGCTGGCTGGTTTATTGCTGATAAATCTGGAGCCGGTGAGCGTGGGTCTCGCGGTATCATTGCAGCACTGGGGCCAGATGGTAAGCCCTCCCGTATCGTAGTTATCTACACGACGGGGAGTCAGGCAACTATGGATGAACGAAATAGACAGATCGCTGAGATAGGTGCCTCACTGATTAAGCATTGGTAACTGTCAGACCAAGTTTACTCATATATACTTTAGATTGATTTAAAACTTCATTTTTAATTTAAAAGGATCTAGGTGAAGATCCTTTTTGATAATCTCATGACCAAAATCCCTTAACGTGAGTTTTCGTTCCACTGAGCGTCAGACCCCGTAGAAAAGATCAAAGGATCTTC

>MjA9Acd-RS.pDule2

TTGAGATCGTTTTGGTCTGCGCGTAATCTCTTGCTCTGAAAACGAAAAAACCGCCTTGCAGGGCGGTTTTTCGAAGGTTCTCTGAGCTACCAACTCTTTGAACCGAGGTAACTGGCTTGGAGGAGCGCAGTCACCAAAACTTGTCCTTTCAGTTTAGCCTTAACCGGCGCATGACTTCAAGACTAACTCCTCTAAATCAATTACCAGTGGCTGCTGCCAGTGGTGCTTTTGCATGTCTTTCCGGGTTGGACTCAAGACGATAGTTACCGGATAAGGCGCAGCGGTCGGACTGAACGGGGGGTTCGTGCATACAGTCCAGCTTGGAGCGAACTGCCTACCCGGAACTGAGTGTCAGGCGTGGAATGAGACAAACGCGGCCATAACAGCGGAATGACACCGGTAAACCGAAAGGCAGGAACAGGAGAGCGCACGAGGGAGCCGCCAGGGGGAAACGCCTGGTATCTTTATAGTCCTGTCGGGTTTCGCCACCACTGATTTGAGCGTCAGATTTCGTGATGCTTGTCAGGGGGGCGGAGCCTATGGAAAAACGGCTTTGCCGCGGCCCTCTCACTTCCCTGTTAAGTATCTTCCTGGCATCTTCCAGGAAATCTCCGCCCCGTTCGTAAGCCATTTCCGCTCGCCGCAGTCGAACGACCGAGCGTAGCGAGTCAGTGAGCGAGGAAGCGGAATATATCCTGTATCACATATTCTGCTGACGCACCGGTGCAGCCTTTTTTCTCCTGCCACATGAAGCACTTCACTGACACCCTCATCAGTGCCAACATAGTAAGCCAGTATACACTCCGCTAGCGCTGATGTCCGGCGGTGCTTTTGCCGTTACGCACCACCCCGTCAGTAGCTGAACAGGAGGGACAGCTCCCGGCGGATTTGTCCTACTCAGGAGAGCGTTCACCGACAAACAACAGATAAAACGAAAGGCCCAGTCTTTCGACTGAGCCTTTCGTTTTATTTGATGCCTGGCAGTTCCCTACTCTCGCATGGGGAGACCCCACACTACCATCGGCGCTACGGCGTTTCACTTCTGAGTTCGGCATGGGGTCAGGTGGGACCACCGCGCTACTGCCGCCAGGCAAATTCTGTTTTATCAGACCGCTTCTGCGTTCTGATTTAATCTGTATCAGGCTGAAAATCTTCTCTCATCCGCCAAAACAGCCAAGCTGGAGACCGTTTAAACTCAATGATGATGATGATGATGGTCGACGGCGCTATTCAGATCCTCTTCTGAGATGAGTTTTTGTTCGGGCCCAAGCTTCGAATTCCCATATGGTACCCGTTTGAAACTGCAGTTATAATCTCTTTCTAATTGGCTCTAAAATCTTTATAAGTTCTTCAGCTACAGCATTTTTTAAATCCATTGGATGCAATTCCTTATTTTTAAATAAACTCTCTAACTCCTCATAGCTATTAACTGTCAAATCTCCACCAAATTTTTCTGGCCTTTTTATGGTTAAAGGATATTCAAGGAAGTATTTAGCTATCTCCATTATTGGATTTCCTTCAACAACTCCAGCTGGGCAGTATGCTTTCTTTATCTTAGCCCTAATCTCTTCTGGAGAGTCATCAACAGCTATAAAATTCCCTTTTGAAGAACTCATCTTTCCTTCTCCATCCAAACCCGTTAAGACAGGGTTGTGAATACAAACAACCTTTTTTGGTAAAAGCTCCCTTGCTAACATGTTTATTTTTCTCTGCTCCATCCCTCCAACTGCAACATCAACGCCAGTATAATGAATAGAATTAACCTGCATTATTGGATAGATAACTTCAGCAACCTTTGGATTTTCATCCTCTCTTGCTATAAGTTCCATACTCCTTCTTGCTCTTTTTAAGGTAGTTTTTAAAGCCAATCTATAGACATTCAGTGTATAATCCTTATCAAGCTCCAGTTCACTTCCATAAACATATTTTGCCTTTAACCCCATTGCTTCAAAAACTTTTTTGTTATAATCTCCTATTTTTCTAATCTCATCCAACTCTCCTTTCTGGTTTAAATAGGCGTGTAAATCAGCCAAATCTATAATTATATCAAATCCAGCATTTTGTAAATCAATCATCTTTTTTATTTGGAGATAATGCCCTAAATGTATTTTACCACTTGGTTCAAAACCTATCGCAGCAGATTTTTCATCTTTTTTTAAAACCTCTCTTAACTCTTCCTCGCTGATAATTTCAGATGTGTTTCTCTTTATCATTTCAAATTCGTCCATGGGGGATTCCTCAAAGCGTAAACTCAGCGTTACAAGTATTACACAAAGTTTTTTATGTTGAGAATATTTTTTTGATGGGGCGCCACTTATTTTTGATCGTTCGCTCAAAGAAGCGGCGCCAGGGTTGTTTTTCTTTTCACCGGTGAGACGGGCAACAGAACGCCATGAGCGGCCTCATTTCTTATTCTGAGTTACAACAGTCCGCACCGCTGTCCGTATATATGAGTAAACTTGGTCCCGGGTTACCGGTTTGGTTAGCGAGAAGAGCCAGTAAAAGACGCAGTGACGGCAATGTCTGATGCAATATGGACAATTGGTTTCTTCTCTGAATGGCGGGAGTATGAAAAGTATGGCTGAAGCGCAAAATGATCCCCTGCTGCCGGGATACTCGTTTAATGCCCATCTGGTGGCGGGTTTAACGCCGATTGAGGCCAACGGTTATCTCGATTTTTTTATCGACCGACCGCTGGGAATGAAAGGTTATATTCTCAATCTCACCATTCGCGGTCAGGGGGTGGTGAAAAATCAGGGACGAGAATTTGTTTGCCGACCGGGTGATATTTTGCTGTTCCCGCCAGGAGAGATTCATCACTACGGTCGTCATCCGGAGGCTCGCGAATGGTATCACCAGTGGGTTTACTTTCGTCCGCGCGCCTACTGGCATGAATGGCTTAACTGGCCGTCAATATTTGCCAATACGGGGTTCTTTCGCCCGGATGAAGCGCACCAGCCGCATTTCAGCGACCTGTTTGGGCAAATCATTAACGCCGGGCAAGGGGAAGGGCGCTATTCGGAGCTGCTGGCGATAAATCTGCTTGAGCAATTGTTACTGCGGCGCATGGAAGCGATTAACGAGTCGCTCCATCCACCGATGGATAATCGGGTACGCGAGGCTTGTCAGTACATCAGCGATCACCTGGCAGACAGCAATTTTGATATCGCCAGCGTCGCACAGCATGTTTGCTTGTCGCCGTCGCGTCTGTCACATCTTTTCCGCCAGCAGTTAGGGATTAGCGTCTTAAGCTGGCGCGAGGACCAACGTATCAGCCAGGCGAAGCTGCTTTTGAGCACCACCCGGATGCCTATCGCCACCGTCGGTCGCAATGTTGGTTTTGACGATCAACTCTATTTCTCGCGGGTATTTAAAAAATGCACCGGGGCCAGCCCGAGCGAGTTCCGTGCCGGTTGTGAAGAAAAAGTGAATGATGTAGCCGTCAAGTTGTCATAATTGGTAACGAATCAGACAATTGACGGCTTGACGGAGTAGCATAGGGTTTGCAGAATCCCTGCTTCGTCCATTTGACAGGCACATTATGCATGCCGCTTCGCCTTCGCGCGCGAATTGATCTGCTGCCTCGCGCGTTTCGGTGATGACGGTGAAAACCTCTGACACATGCAGCTCCCGGAGACGGTCACAGCTTGTCTGTAAGCGGATGCCGGGAGCAGACAAGCCCGTCAGGGCGCGTCAGCGGGTGTTGGCGGGTGTCGGGGCGCAGCCATGACCCAGTCAACTGCGATGAGTGGCAGGGCGGGGCGTAATTTTTTTAAGGCAGTTATTGGTGCCCTTAAACGCCTGGTTGCTACGCCTGAATAAGTGATAATAAGCGGATGAATGGCAGAAATTCGAAAGCAAATTCGACCCGGTCGTCGGTTCAGGGCAGGGTCGTTAAATAGCCGCTTATGTCTATTGCTGGTTTACCGGTTTATTGACTACCGGAAGCAGTGTGACCGTGTGCTTCTCAAATGCCTGAGGCCAGTTTGCTCAGGCTCTCCCCGTGGAGGTAATAATTGACGATATGATCATTTATTCTGCCTCCCAGAGCATGATAAAAACGGTTAGCGCTTCGTTAATACAGATGTAGGTGTTCCACAGGGTAGCCAGCAGCATCCTGCGATGCAGATCCGGAACATAATGGTGCAGGGCGCTTGTTTCGGCGTGGGTATGGTGGCAGGCCCCGTGGCCGGGGGACTGTTGGGCGCTGCCGGCACCTGTCCTACGAGTTGCATGATAAAGAAGACAGTCATAAGTGCGGCGACGATAGTCATGCCCCGCGCCCACCGGAAGGAGCTACCGGCAGCGGTGCGGACTGTTGTAACTCAGAATAAGAAATGAGGCCGCTCATGGCGTTCTGTTGCCCGTCTCACTGGTGAAAAGAAAAACAACCCTGGCGCCGCTTCTTTGAGCGAACGATCAAAAATAAGTGGCGCCCCATCAAAAAAATATTCTCAACATAAAAAACTTTGTGTAATACTTGTAACGCTGAATTCCCGGCGGTAGTTCAGCAGGGCAGAACGGCGGACTCTAAATCCGCATGGCGCTGGTTCAAATCCGGCCCGCCGGACCACTGCAGATCCTTAGCGAAAGCTAAGGATTTTTTTTAAGCTTGGCACTGGCCGTCGTTTTACAACGTCGTGACTGGGAAAACCCTGGCGTTACCCAACTTAATCGCCTTGCAGCACATCCCCCTTTCGCCAGACGCTCTCCCTTATGCGACTCCTGCATTAGGAAGCAGCCCAGTAGTAGGTTGAGGCCGTTGAGCACCGCCGCCGCAAGGAATGGTGCATGCAAGGAGCCCGAGATGCGCCGCGTGCGGCTGCTGGAGATGGCGGACGCGATGGATATGTTCTGCCAAGGGTTGGTTTGCGCATTCACAGTTCTCCGCAAGAATTGATTGGCTCCAATTCTTGGAGTGGTGAATCCGTTAGCGAGGTGCCGCCGGCTTCCATTCAGGTCGAGGTGGCCCGGCTCCATGCACCGCGACGCAACGCGGGGAGGCAGACAAGGTATAGGGCGGCGCCTACAATCCATGCCAACCCGTTCCATGTGCTCGCCGAGGCGGCATAAATCGCCGTGACGATCAGCGGTCCAATGATCGAAGTTAGGCTGGTAAGAGCCGCGAGCGATCCTTGAAGCTGTCCCTGATGGTCGTCATCTACCTGCCTGGACAGCATGGCCTGCAACGCGGGCATCCCGATGCCGCCGGAAGCGAGAAGAATCATAATGGGGAAGGCCATCCAGCCTCGCGTCGCGAACGCCAGCAAGACGTAGCCCAGCGCGTCGGCCGCCATGCCGGCGATAATGGCCTGCTTCTCGCCGAAACGTTTGGTGGCGGGACCAGTGACGAAGGCTTGAGCGAGGGCGTGCAAGATTCCGAATACCGCAAGCGACAGGCCGATCATCGTCGCGCTCCAGCGAAAGCGGTCCTCGCCGAAAATGACCCAGAGCGCTGCCGGCACCTGTCCTACGAGTTGCATGATAAAGAAGACAGTCATAAGTGCGGCGACGATAGTCATGCCCCGCGCCCACCGGAAGGAGCTGACTGGGTTGAAGGCTCTCAAGGGCATCGGTCGACGCTCTCCCTTATGCGACTCCTGCATTAGGAAGCAGCCCAGTAGTAGGTTGAGGCCGTTGAGCACCGCCGCCGCAAGGAATGGTGCATGCAAGGAGATGGCGCCCAACAGTCCCCCGGCCACGGGGCCTGCCACCATACCCACGCCGAAACAAGCGCTCATGAGCCCGAAGTGGCGAGCCCGATCTTCCCCATCGGTGATGTCGGCGATATAGGCGCCAGCAACCGCACCTGTGGCGCCGGTGATGCCGGCCACGATGCGTCCGGCGTAGAGGATCCCGGTAAACCAGCAATAGACATAAGCGGCTATTTAACGACCCTGCCCTGAACCGACGACCGGGTCATCGTGGCCGGATCTTGCGGCCCCTCGGCTTGAACGAATTGTTAGACATTATTTGCCGACTACCTTGGTGATCTCGCCTTTCACGTAGTGGACAAATTCTTCCAACTGATCTGCGCGCGAGGCCAAGCGATCTTCTTCTTGTCCAAGATAAGCCTGTCTAGCTTCAAGTATGACGGGCTGATACTGGGCCGGCAGGCGCTCCATTGCCCAGTCGGCAGCGACATCCTTCGGCGCGATTTTGCCGGTTACTGCGCTGTACCAAATGCGGGACAACGTAAGCACTACATTTCGCTCATCGCCAGCCCAGTCGGGCGGCGAGTTCCATAGCGTTAAGGTTTCATTTAGCGCCTCAAATAGATCCTGTTCAGGAACCGGATCAAAGAGTTCCTCCGCCGCTGGACCTACCAAGGCAACGCTATGTTCTCTTGCTTTTGTCAGCAAGATAGCCAGATCAATGTCGATCGTGGCTGGCTCGAAGATACCTGCAAGAATGTCATTGCGCTGCCATTCTCCAAATTGCAGTTCGCGCTTAGCTGGATAACGCCACGGAATGATGTCGTCGTGCACAACAATGGTGACTTCTACAGCGCGGAGAATCTCGCTCTCTCCAGGGGAAGCCGAAGTTTCCAAAAGGTCGTTGATCAAAGCTCGCCGCGTTGTTTCATCAAGCCTTACGGTCACCGTAACCAGCAAATCAATATCACTGTGTGGCTTCAGGCCGCCATCCACTGCGGAGCCGTACAAATGTACGGCCAGCAACGTCGGTTCGAGATGGCGCTCGATGACGCCAACTACCTCTGATAGTTGAGTCGATACTTCGGCGATCACCGCTTCCCTCATACTCTTCCTTTTTCAATATTATTGAAGCATTTATCAGGGTTATTGTCTCATGAGCGGATACATATTTGAATGTATTTAGAAAAATAAACAAATAGCTAGCTCACTCGGTCGCATCGATGATAAGCTGTCAAACATGAGAATTACAACTTATATCGTATGGGGCTGACTTCAGGTGCTACATTTGAAGAGATAAATTGCACTGAAATCTAGAAATATTTTATCTGATTAATAAGATGATCTTC

## Supplementary Figures

**Figure S1.**
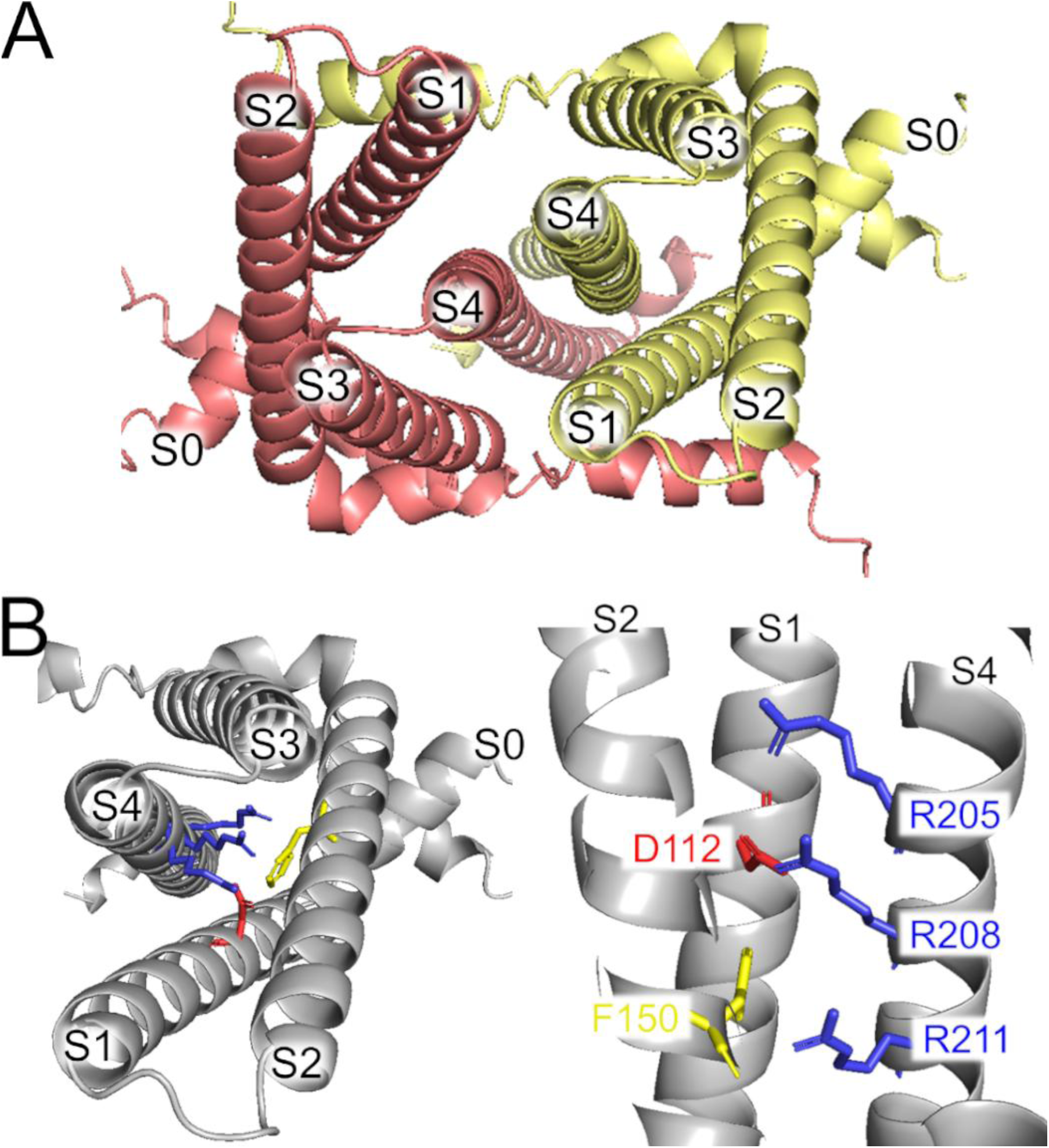
AlphaFold dimer hHv1 structural model. (A) Extracellular view of the model, showing each subunit in a different color. (B) One subunit of the dimer showing the position of the arginine gating charges in S4 (blue), the selectivity filter (red), and the charge transfer center (yellow).

**Figure S2.**
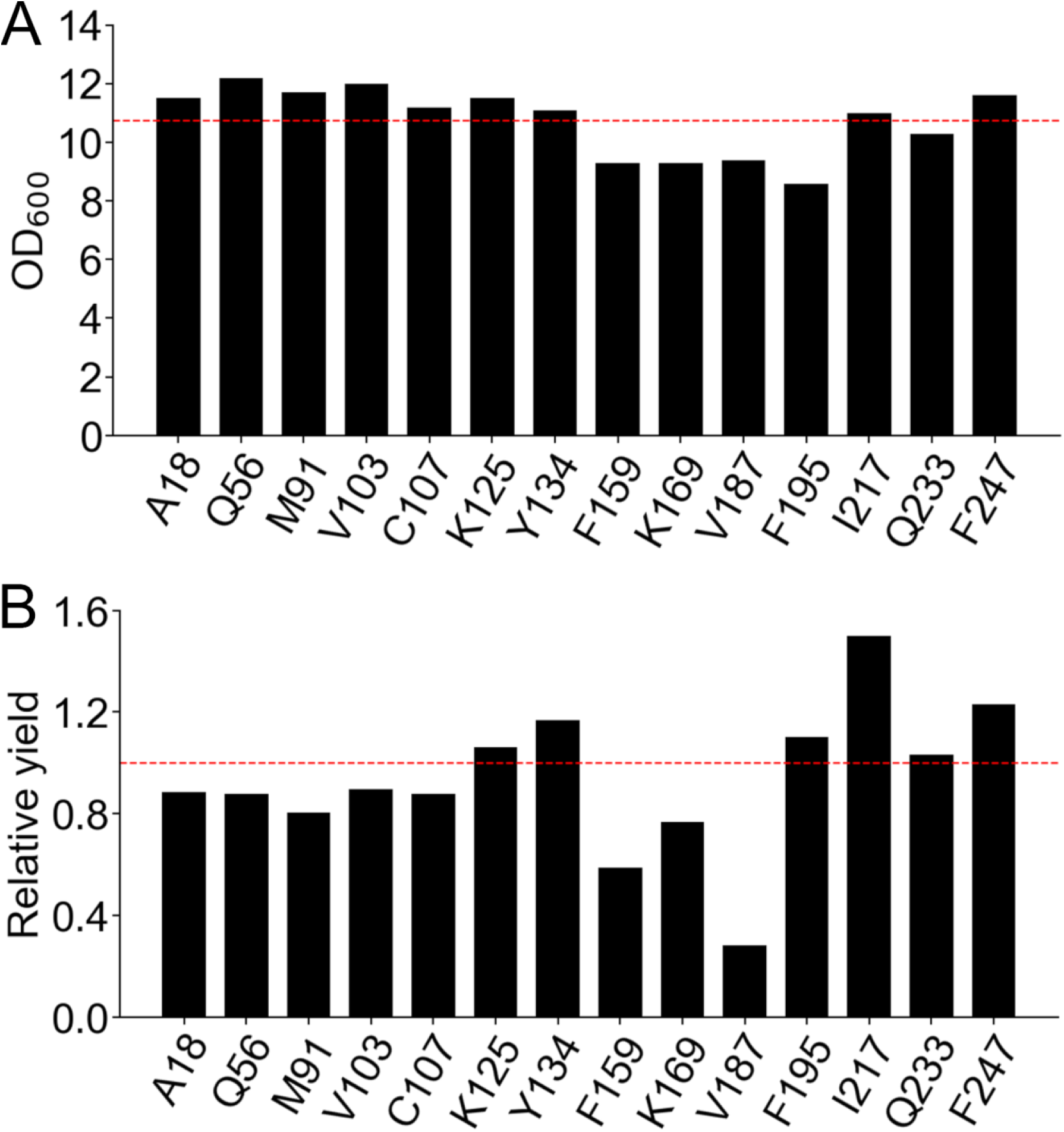
The protein yield of hH_v_1-Acd depended on the position of Acd incorporation. (A) Optical density at 600 nm (OD_600_) of the final cultures expressing the hH_v_1 with Acd replacing the indicated amino acid. (B) Relative protein yield calculated by multiplying the OD_600_ and Western blot band intensity of cellular extracts from the final cultures expressing the hH_v_1 with Acd replacing the indicated amino acid. Values of normalized by the No TAG condition, which are cells expressing the hH_v_1 without any amber stop codon in the presence of Acd and the aminoacyl-tRNA synthetase/tRNA pair (red broken line).

**Figure S3.**
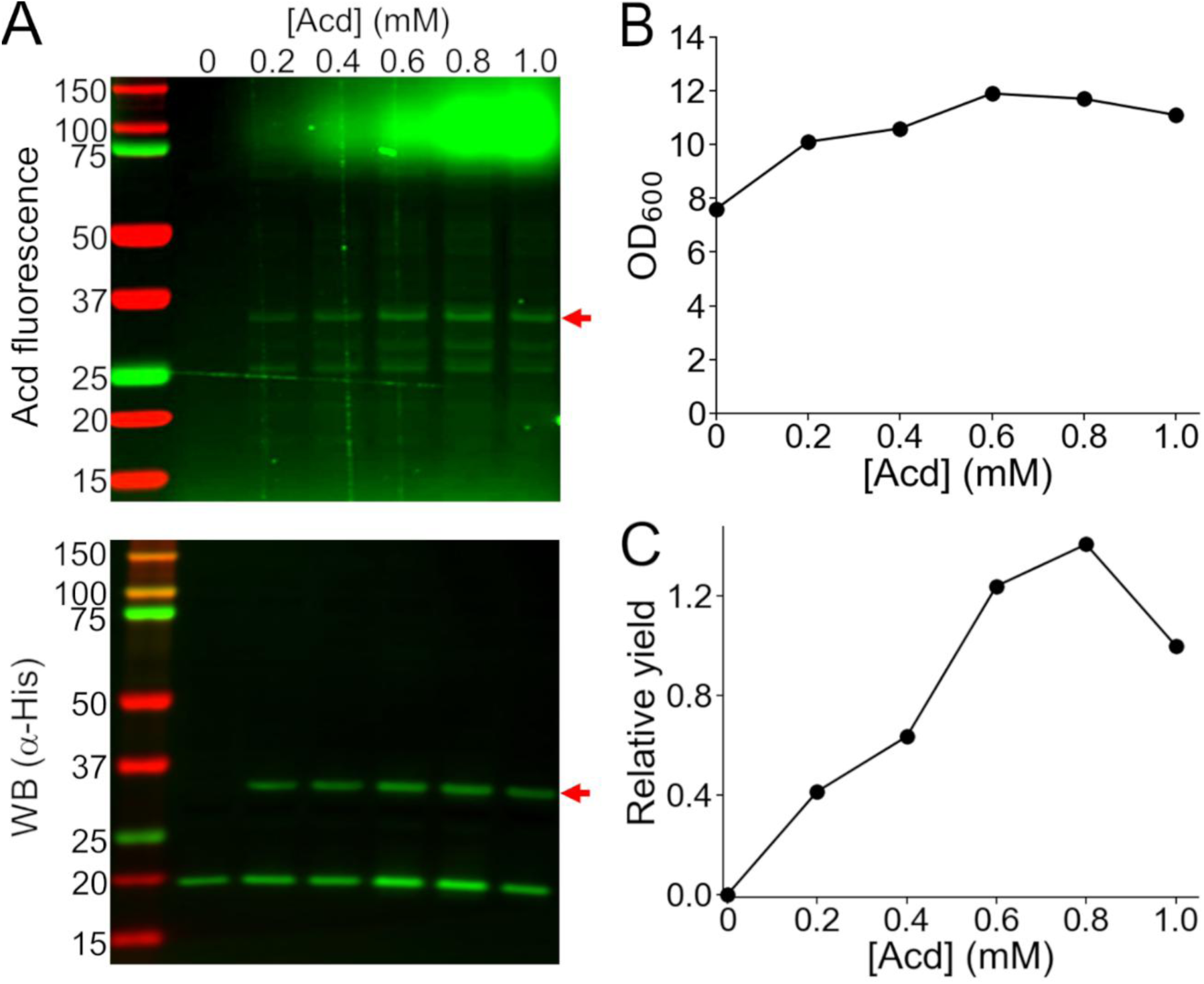
The truncation of hH_v_1-K125Acd was not decreased at higher concentrations of Acd in the culture medium. (A) Cellular extracts of cells co-transformed with the MjA9-AcdRS/tRNA pair plasmid and hH_v_1-K125TAG and grown with the indicated concentration of Acd in the culture medium were separated by SDS-PAGE to visualize the hH_v_1 band (red arrow) by Acd fluorescence (top) and Western blot against the N-terminus His-tag (bottom). (B) Optical density at 600 nm (OD_600_) of the cultures at the end of the expression. (C) Relative yield obtained by multiplying the Western blot band intensity by the OD600, normalized by the value obtained at 1 mM Acd.

**Figure S4.**
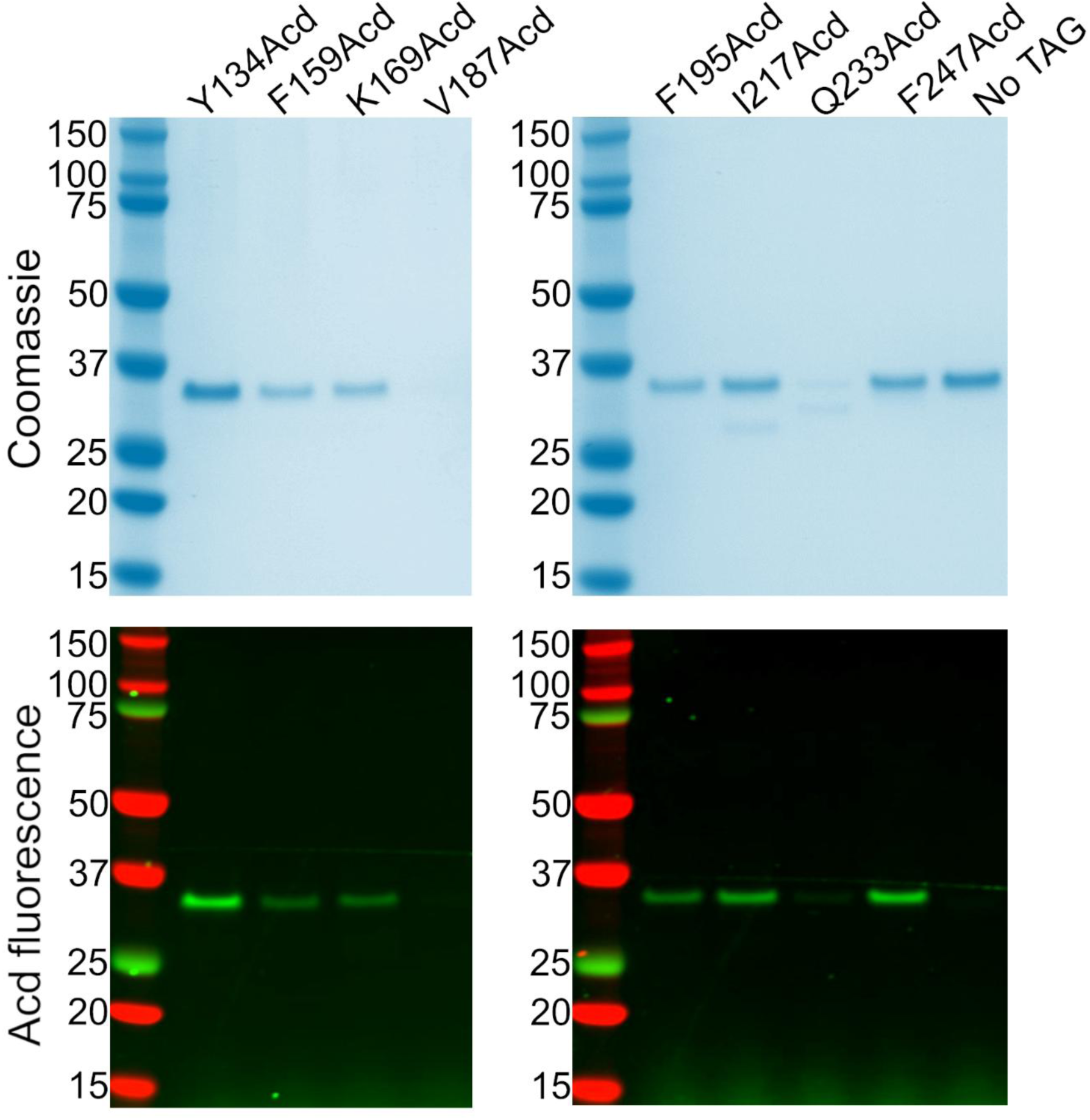
SDS-PAGE of the hH_v_1 protein samples after purification. Coomassie-stained (top) and Acd fluorescence (bottom) gels of the samples from the elution of the nickel resin column, separated by SDS-PAGE. Note that V187Acd and Q233Acd did not contain appreciable amounts of protein. The No TAG condition consisted of samples purified from cells expressing the hH_v_1 without any amber stop codon in the presence of Acd and the aminoacyl-tRNA synthetase/tRNA pair.

**Figure S5.**
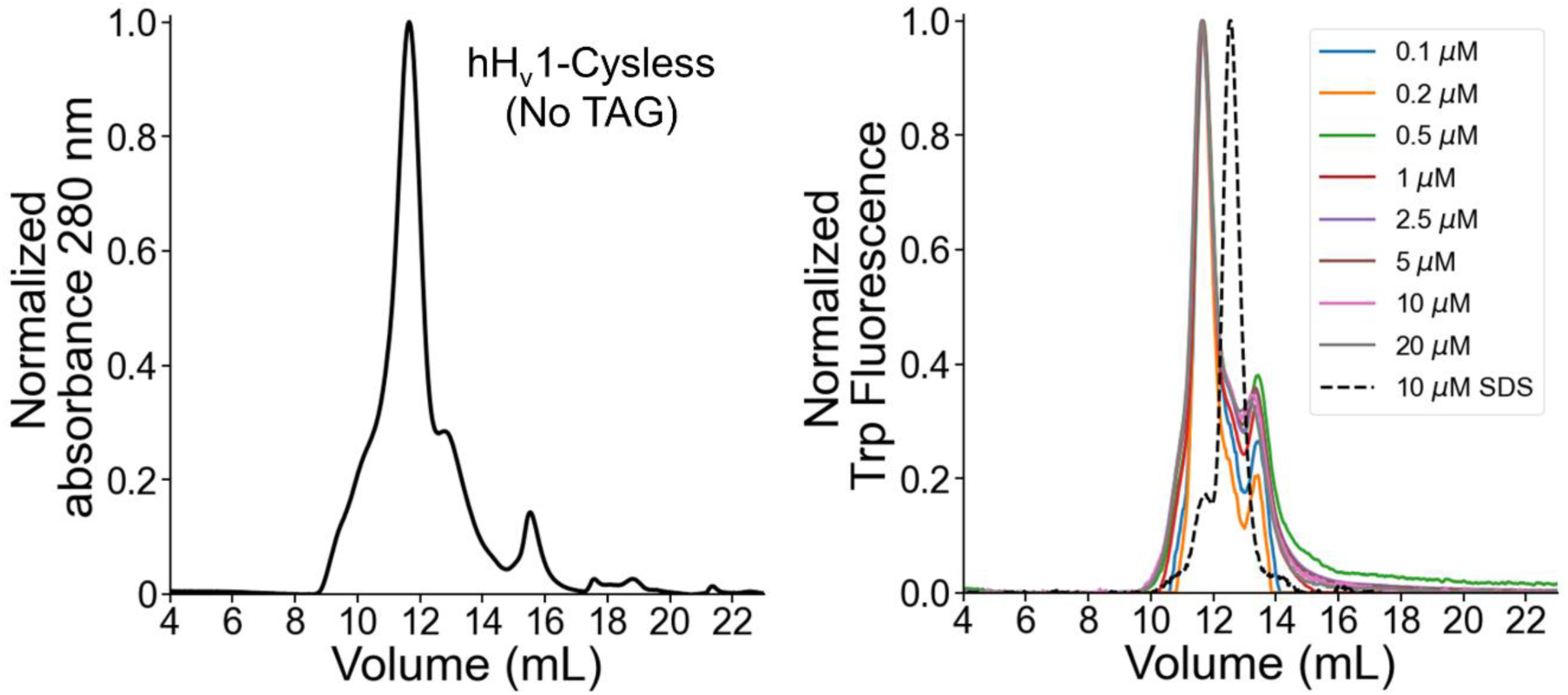
The main hH_v_1 peak in the size exclusion chromatogram corresponds to the dimer. (Left) The No TAG sample was purified from cells expressing the hH_v_1 without any amber stop codon in the presence of Acd and the aminoacyl-tRNA synthetase/tRNA pair. The sample eluted from the immobilized affinity chromatography column was concentrated and injected into the size exclusion chromatography column. (Right) SEC-cleaned samples were injected at the indicated concentration. The elution volumes of the samples were similar, with a mean ± standard deviation of 11.64 ± 0.01 mL. Additionally, a sample was run in a denaturing buffer containing 50 mM Tris, pH 8.0, 150 mM NaCl, and 0.2% SDS (dashed lines; elution volume of 12.5 mL). The invariant elution volume at low concentrations and higher elution volume in denaturing buffer strongly suggests that the main peak on the hH_v_1 chromatograms corresponds to the dimer. The delay between the fluorescence and absorbance peaks is due to the physical separation of the detectors.

**Figure S6.**
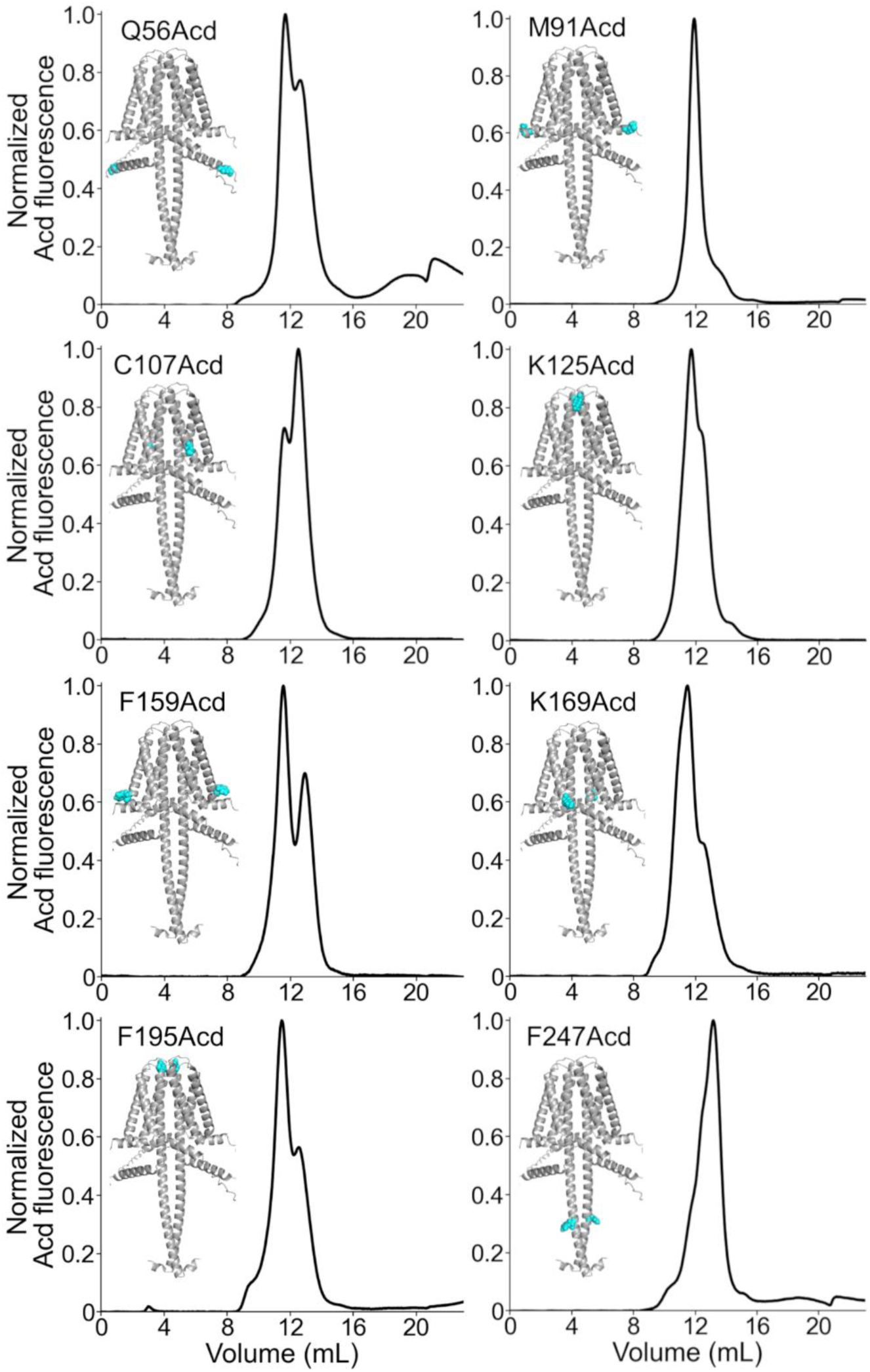
Fluorescence-detection size exclusion chromatography of hH_v_1-Acd proteins. Chromatograms of the purified hH_v_1 proteins with Acd incorporated at the indicated amino acid position. The AlphaFold dimer model with the amino acid replaced by Acd as cyan spheres is included.

**Figure S7.**
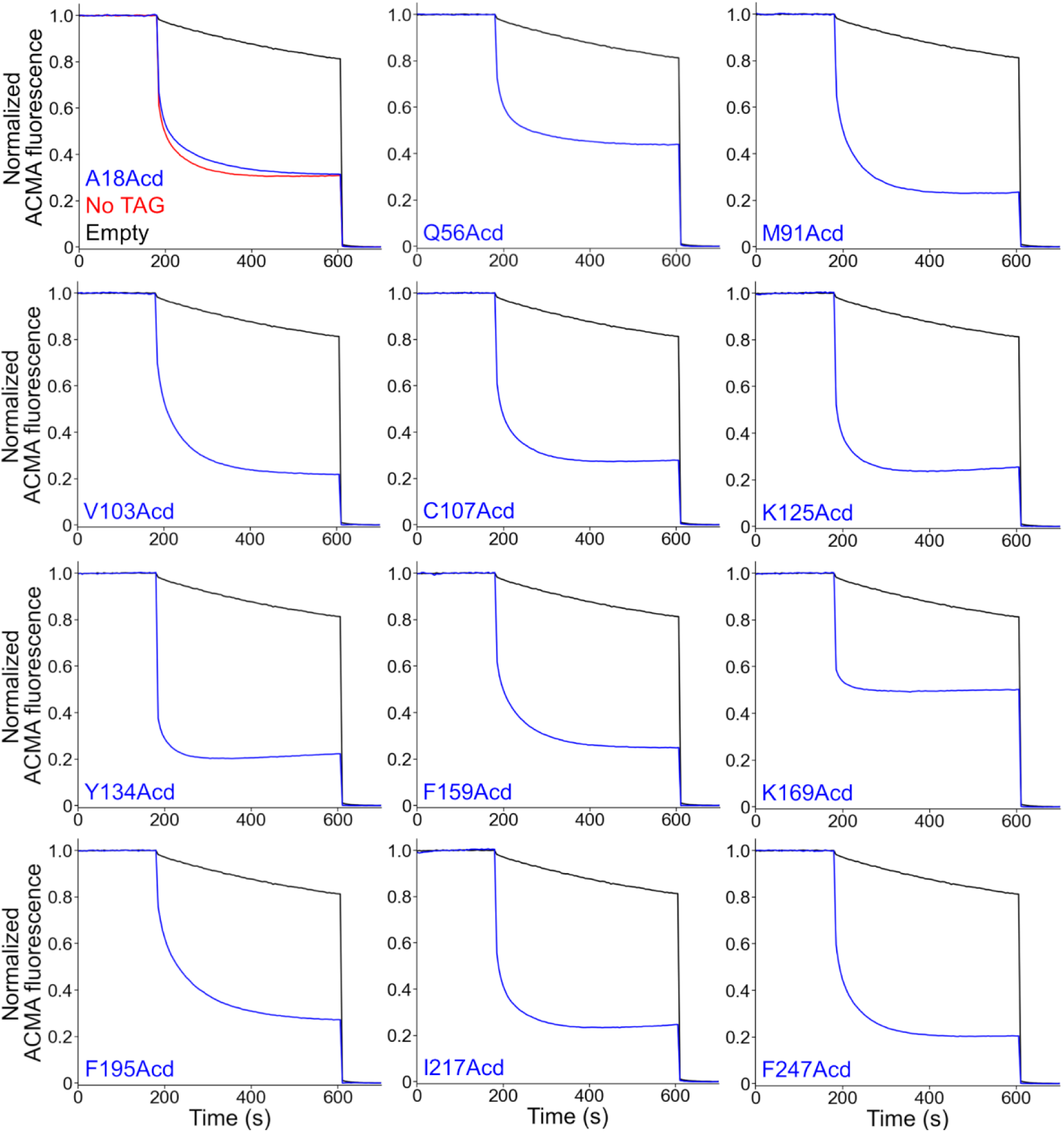
The purified hH_v_1-Acd proteins were functional proton channels. Liposome proton flux assays of asolectin proteoliposomes containing the indicated hH_v_1-Acd protein. The proteoliposome samples produced ACMA fluorescence quenching after the addition of valinomycin (180 s) in the presence of a potassium gradient. The protonophore CCCP was added at the end of the experiment (600 s). The No TAG sample (red trace) contains proteoliposomes containing hH_v_1 without any amber stop codon expressed in the presence of Acd and the aminoacyl-tRNA synthetase/tRNA pair. Empty asolectin liposomes (black traces) showed slow ACMA fluorescence quenching.

**Fig. S8.**
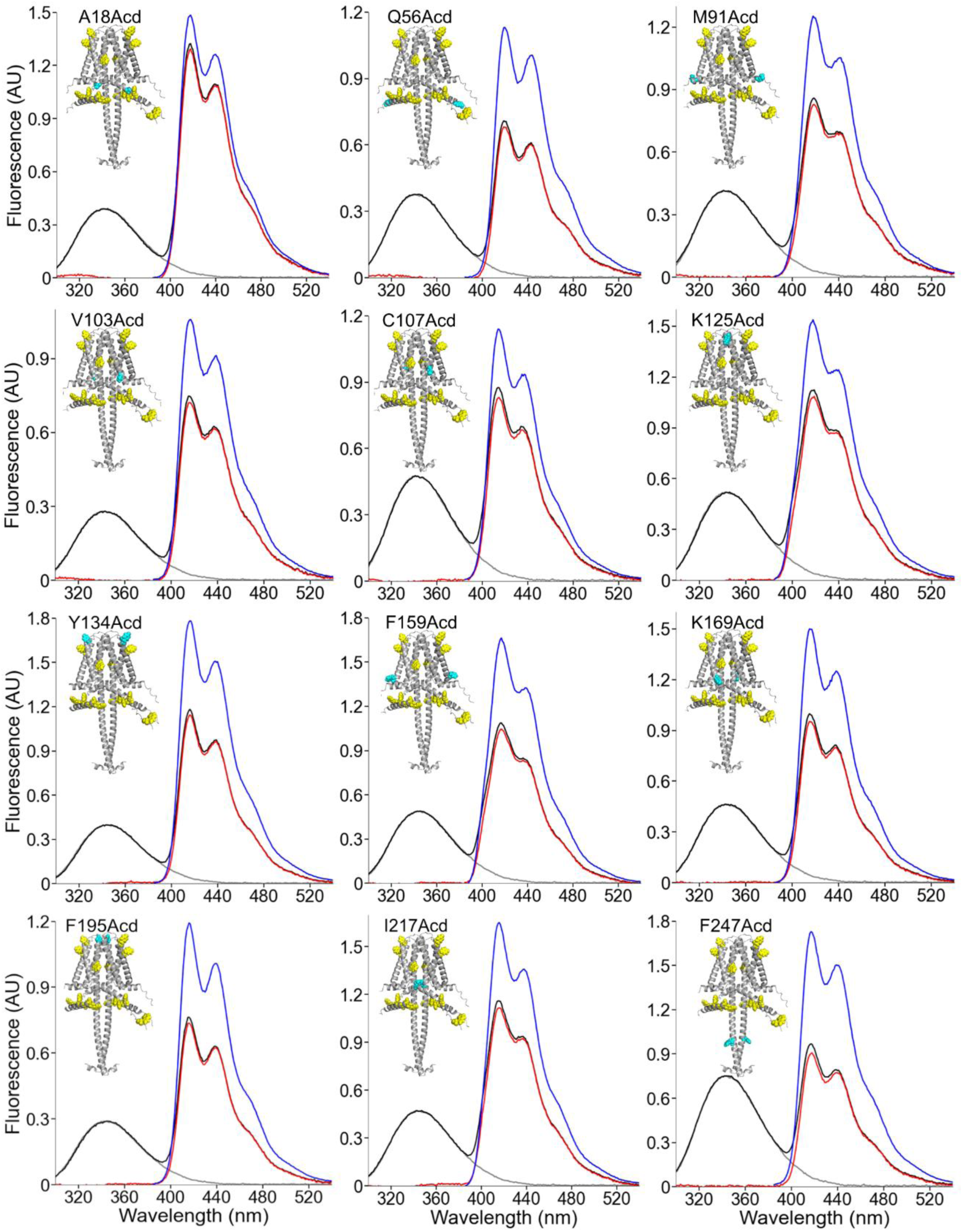
Fluorescence emission spectra of hH_v_1-Acd. Spectra of the indicated hH_v_1-Acd protein sample obtained when exciting at 280 nm (black) and 370 nm (blue). The isolated spectrum of Acd (red) was obtained by subtracting the black trace from the normalized spectrum of the No TAG sample excited at 280 nm (grey).

**Figure S9.**
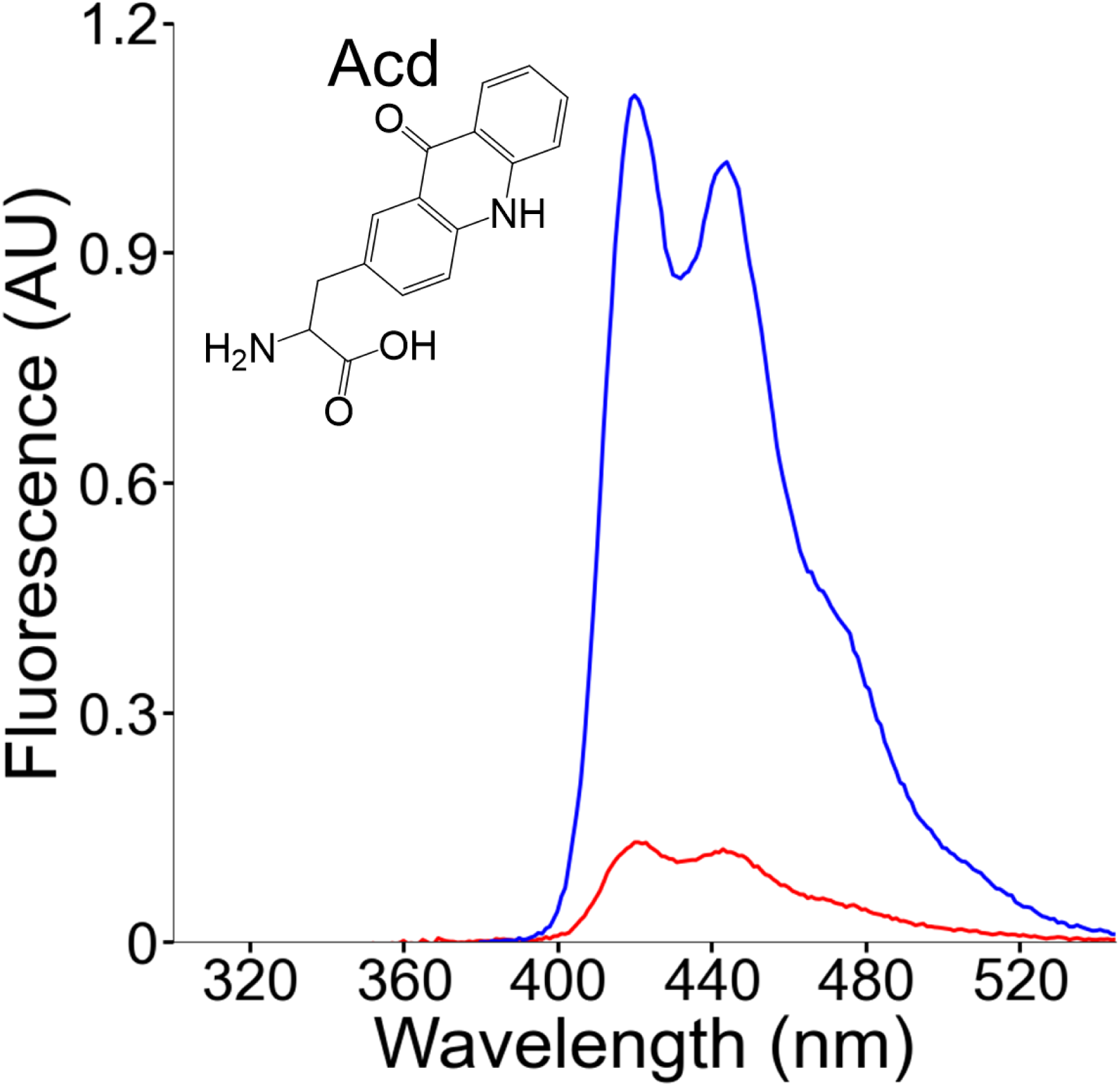
Fluorescence emission spectra of the Acd amino acid in solution. Spectra of Acd in Buffer-H2 using an excitation wavelength of 370 nm (blue) or 280 nm (red).

**Figure S10.**
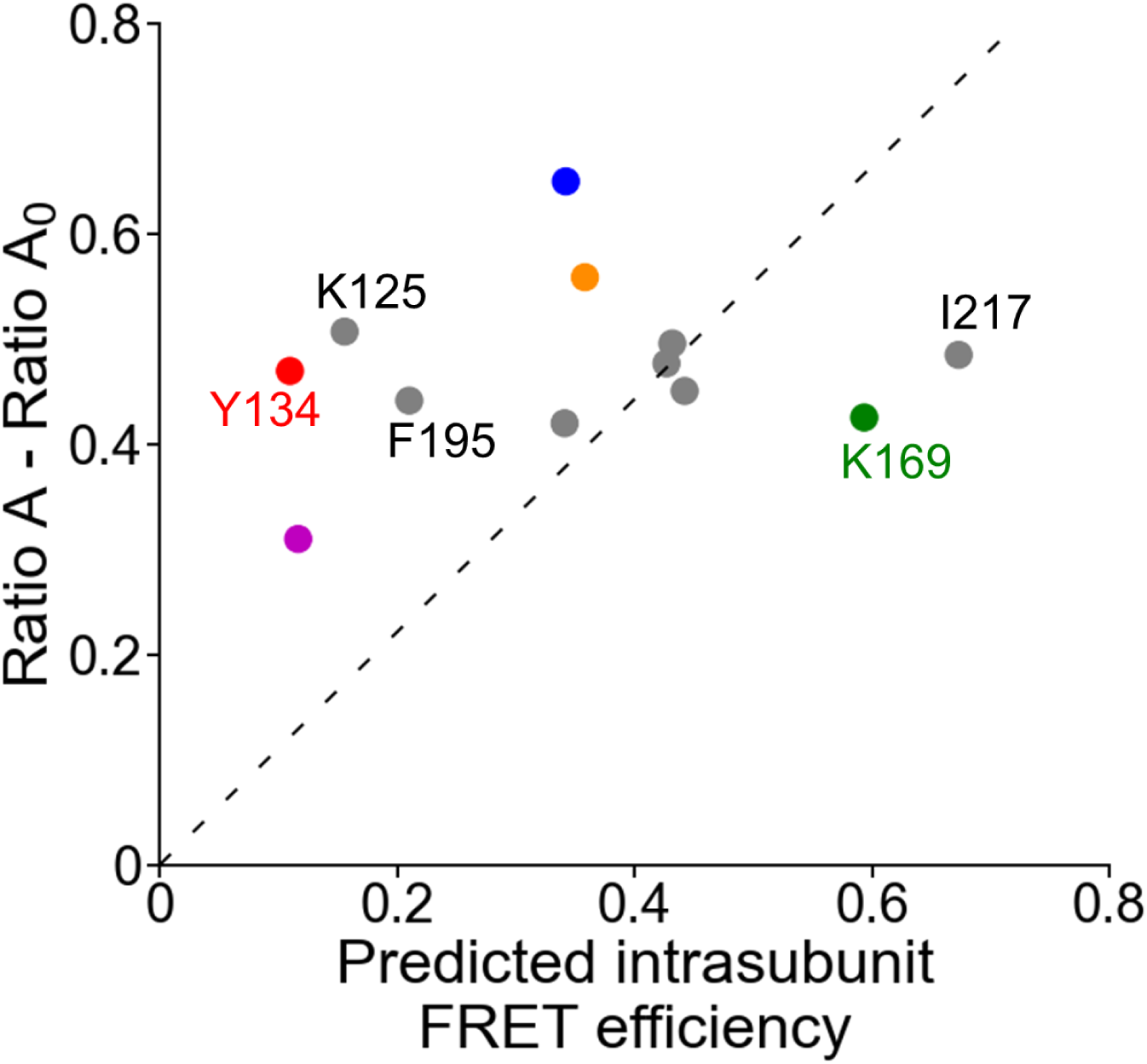
Fluorescence emission spectra of the Acd amino acid in solution. Mean *Ratio A – Ratio A_0_* values (410-480 nm; N=4) as a function of the predicted intrasubunit FRET efficiencies from one subunit of the AlphaFold hH_v_1 structural model, colored according to Figure 5F. The broken line is the best fit to the equation (*Ratio A – Ratio A_0_*) = m(Predicted intrasubunit FRET efficiency).

**Fig. S11.**
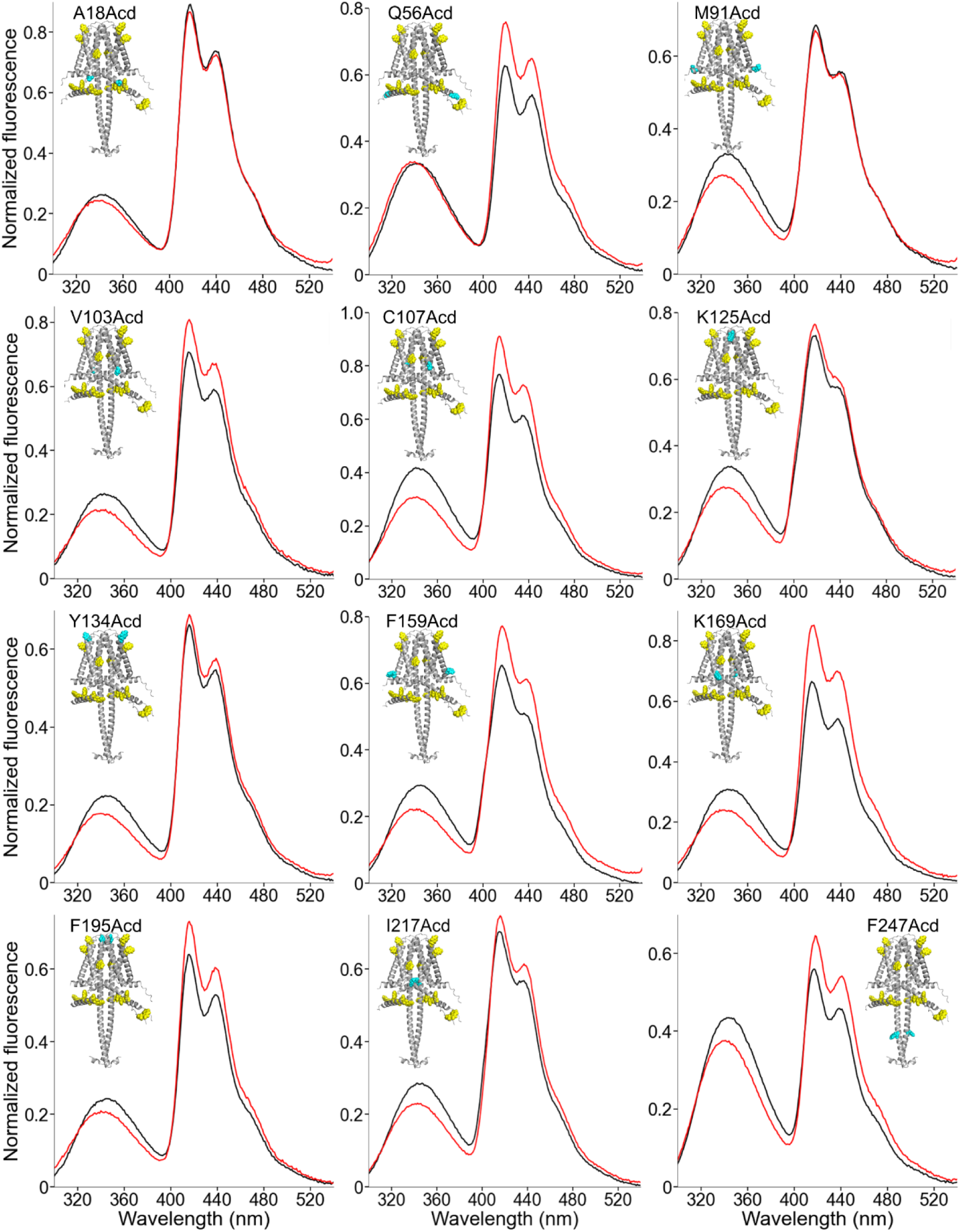
Changes in the fluorescence emission spectra of hH_v_1-Acd in the presence of zinc. Spectra of the indicated hH_v_1-Acd protein sample obtained when exciting at 280 nm in the Apo state (black) and in the presence of 1 mM Zn^2+^ (red). Spectra were normalized by the maximum intensity of the Acd emission spectrum of the same sample excited at 370 nm.

**Figure S12.**
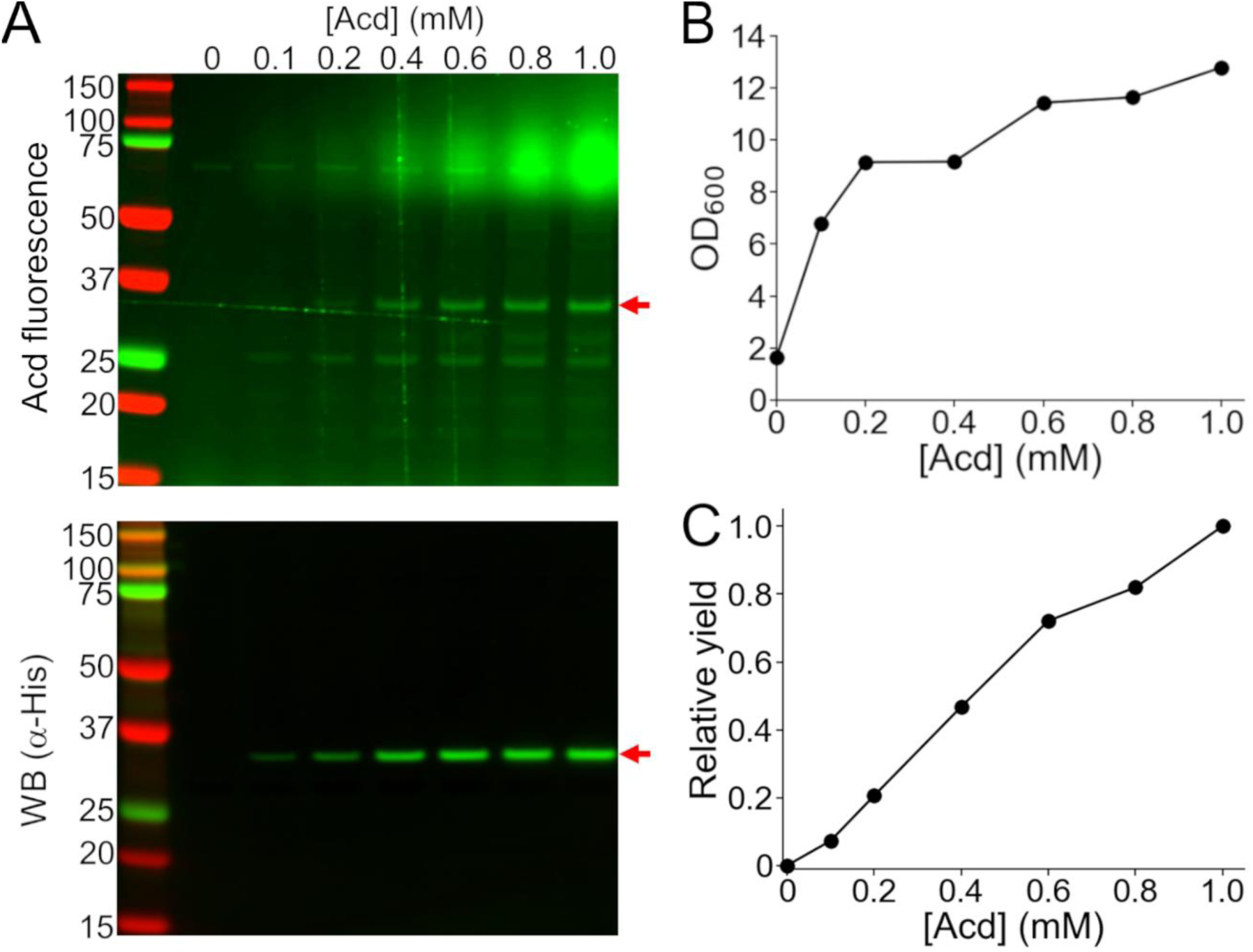
The protein yield of hH_v_1-A18Acd was proportional to the concentration of Acd in the culture medium. (A) Cellular extracts of cells co-transformed with the MjA9-AcdRS/tRNA pair plasmid and hH_v_1-A18TAG and grown with the indicated concentration of Acd in the culture medium were separated by SDS-PAGE to visualize the hH_v_1 band (red arrow) by Acd fluorescence (top) and Western blot against the N-terminus His-tag (bottom). (B) Optical density at 600 nm (OD_600_) of the cultures at the end of the expression. (C) Relative yield obtained by multiplying the Western blot band intensity by the OD_600_, normalized by the value obtained at 1 mM Acd.

## Supplementary Tables

**Table S1.** Size-exclusion chromatography elution volumes of the hH_v_1 proteins. Elution volumes were measured as the location of the maximum value of absorbance at 280 nm in the chromatograms.

| hHv1 sample | Elution volume (mL) |
| --- | --- |
| No TAG | 11.5 |
| A18Acd | 11.6 |
| Q56Acd | 11.7 |
| M91Acd | 11.9 |
| V103Acd | 11.5 |
| C107Acd | 12.5 |
| K125Acd | 11.7 |
| Y134Acd | 11.5 |
| F159Acd | 11.5 |
| K169Acd | 11.5 |
| F195 Acd | 11.5 |
| I217Acd | 12.4 |
| F247Acd | 13.1 |

**Table S2.** Emission fluorescence spectrum maximum of Acd in different solvents. The wavelength corresponds to the location of the maximum intensity of the emission fluorescence spectrum of the free amino acid dissolved in the corresponding solvent. EtAc: ethyl acetate, OctOH: 1-octanol, ButOH: 1-butanol, EtOH: ethanol.

| Solvent | Emission Maximum (nm) |
| --- | --- |
| EtAc | 402 |
| OctOH | 414 |
| ButOH | 415 |
| EtOH | 414 |
| Water | 421 |
| Buffer-H2 | 420 |

**Table S3.** The emission fluorescence spectrum maximum of the hH_v_1-Acd proteins. The wavelength corresponds to the location of the maximum intensity of the emission fluorescence spectrum of the samples.

| hHv1 sample | Emission Maximum (nm) |
| --- | --- |
| A18Acd | 418 |
| Q56Acd | 420 |
| M91Acd | 419 |
| V103Acd | 417 |
| C107Acd | 415 |
| K125Acd | 418 |
| Y134Acd | 416 |
| F159Acd | 418 |
| K169Acd | 415 |
| F195 Acd | 416 |
| I217Acd | 415 |
| F247Acd | 418 |

---

**Supplementary Dataset S1 (separate file).** Coordinates of the AlphaFold predicted model of the hH_v_1 dimer (in PDB format).

